# Chromosomal assembly of the flat oyster (*Ostrea edulis* L.) genome as a new genetic ressource for aquaculture

**DOI:** 10.1101/2022.06.26.497643

**Authors:** Isabelle Boutet, Homère J. Alves Monteiro, Lyam Baudry, Takeshi Takeuchi, Eric Bonnivard, Bernard Billoud, Sarah Farhat, Ricardo Gonzales-Haraya, Benoit Salaun, Ann Andersen, Jean-Yves Toullec, François Lallier, Jean-François Flot, Nadège Guiglielmoni, Ximing Guo, Li Cui, Bassem Allam, Emmanuelle Pales-Espinoza, Jakob Hemmer-Hansen, Martial Marbouty, Romain Koszul, Arnaud Tanguy

**Affiliations:** Sorbonne Université, CNRS, UMR 7144, Station Biologique de Roscoff, Place Georges Teissier, CS90074, 29688 Roscoff, France; National Institute of Aquatic Resources, Technical University of Denmark, Vejlsøvej 39, 8600, Silkeborg, Denmark; Institut Pasteur, Unité Régulation Spatiale des Génomes, CNRS, Paris, France; Marine Genomics Unit, Okinawa Institute of Science and Technology Graduate University, Onna, Okinawa, Japan; Sorbonne Université, CNRS, UMR 8227, Station Biologique de Roscoff, Place Georges Teissier, CS90074, 29688 Roscoff, France; Marine Animal Disease Laboratory, School of Marine and Atmospheric Sciences, 100 Nicolls Road, Stony Brook University, Stony Brook, NY 11794-5000, USA; Centre Régional de la Conchyliculture Bretagne Nord, 2 Rue du Parc au Duc, 29600 Morlaix, France; Université Libre de Bruxelles, Evolutionary Biology and Ecology, Avenue F.D. Roosevelt 50, B-1050 Brussels, Belgium; Haskin Shellfish Research Laboratory, Department of Marine and Coastal Sciences, Rutgers University, Port Norris, NJ, USA; Department of Marine Organism Taxonomy and Phylogeny, Institute of Oceanology, Chinese Academy of Sciences, Qingdao, Shandong 266071, China

**Keywords:** flat oyster, genome, transposable elements, Martelia, population, aquaculture

## Abstract

The European flat oyster (*Ostrea edulis* L.) is the endemic species of the European coasts. Its exploitation has been reduced during the last decades, because of the appearance of two parasites that have led to the collapse of the stocks and the strong decline of the natural oyster beds. *O. edulis* has been the subject of numerous studies and programs in population genetics and on the presence of the parasites *Bonamia ostreae* and *Marteilia refringens*. These studies investigated the effects of these parasites mainly on immunity at the molecular and cellular levels. Several genetic selection programs especially related to resistance to the parasite have been initiated. Within the framework of a European project (PERLE 2) which aims to produce genetic lines of *O. edulis* with hardiness traits (growth, survival, resistance) for the purpose of repopulating natural oyster beds in Brittany and reviving the culture of this species on the foreshore, obtaining a reference genome has proved to be essential as done recently in many bivalve species of aquaculture interest. Here, we present a chromosome-level genome assembly and annotation for the European flat oyster, generated by combining PacBio technology, Illumina, 10X linked and Hi-C sequencing. The finished assembly is 887.2 Mb with a scaffold-N50 of 97.1 Mb scaffolded on the expected 10 pseudo-chromosomes. Annotation of the genome revealed the presence of 35962 protein-coding genes. We analyzed in details the transposable elements (TE) diversity in the flat oyster genome, highlight some specificities in tRNA and miRNA composition and provide first insights in the molecular response of *O. edulis* to *M. refringens*. This genome will serve as a reference for genomic studies on *O. edulis* to better understand its basic physiology or developing genetic markers in breeding projects for aquaculture or natural reef restoration.

## Background

The European flat oyster, *Ostrea edulis* (NCBI:txid37623), is the native oyster species in Europe and its distribution ranges from the Norwegian Sea in the north to Morocco in the south, and east through the Mediterranean Sea to the Black Sea (OSPAR, 2009). The flat oyster industry has been in decline since the 19th century mainly due to its habitat destruction, over-exploitation (Beck et al., 2011), irregular recruitment and apparition of the parasites *Marteilia refringens* and *Bonamia ostreae* during the 1970s (Smaal et al., 2015). In France, the *O. edulis* the production represents 1100 tons per year and is mainly located in Brittany. Because of its economic and ecological importance, *O. edulis* is subjected to extensive protection and restoration and has been identified as a priority species by the OSPAR Convention (OSPAR, 2009). Indeed, the flat oyster ecosystem is similar to calcareous biogenic reefs and contributes to the substrate stabilization, good water quality and habitat formation for other species (Beck et al., 2011). Unlike the Crassostreides oysters *Crassostrea gigas* or *Crassostrea. virginica, O. edulis* is a larviparous species with an asynchronous hermaphrodite with rhythmic consecutive sexuality. Molecular studies on the physiology of flat oyster, including basics traits and adaptative response to abiotic parameters, are rare and mainly focused on the study of the response of *O. edulis* to its parasite *B. ostreae* mainly in laboratory experiments (Ronza et al., 2018; Gervais et al., 2019; Cocci et al., 2020).

Aquaculture genomics aims to identify the genetic basis of performance and production traits for selective breeding programs(Bernatchez et al., 2017; Shen & Yue, 2019) such as growth in the Atlantic salmon (*Salmo salar*), the Nile tilapia (*Oreochromis niloticus*) or the Pacific white shrimp (*Litopenaeus vannamei*) (Wang et al., 2017). Other traits such as disease resistance, feed conversion efficiency, environmental tolerance, and product quality are also significant but harder to study because of their low heritability (Yáñez et al., 2015) and their late appearance in the life of organisms. Due to the strong impact of the *B. ostreae* parasite on the survival of *O. edulis*, several programs that aimed to identify QTL associated with parasite resistance have been implemented using comparative transcriptomic approaches (Pardo et al., 2016; Ronza et al., 2018) and SNP genotyping (Vera et al., 2019). To partly resolve these limitations and provide accurate markers for aquaculture, sequencing of bivalve genomes has been initiated, including the oysters *C. gigas* (Peñaloza et al., 2021), *C. honkongensis* (Peng et al., 2020), *C. ariakensis* (Li et al., 2021), *C. virginica* (Modak et al., 2021) and *Pinctada fucata* (Du et al., 2017), the scallop *Pecten maximus* (Kenny et al., 2020), the blood clam *Scapharca broughtonii* (Bai et al., 2019), the hard-shelled mussel *Mytilus coruscus* (Yang et al., 2021) and the hard clam *Mercenaria mercenaria* (Song et al., 2021; Farhat et al., 2022). Since 2018, we initiated a selective breeding program (European project PERLE 2) to characterize rusticity parameters, including survival, growth and resistance to parasites in the flat oyster *O. edulis*. The main objectives of this project are (1) the production of a large set of biparental oyster families tested in different environments (deep water and foreshore) and (2) to further contribute to restore natural oyster beds. Furthermore, the generation of genomic tools is essential to address fundamental questions related to evolution, environmental adaptations or in the particular case of *O. edulis* the understanding of hermaphrodism. In the present study, an improved chromosome-level assembly of *O. edulis* genome was developed through a combination of high coverage Pacific Biosciences (PacBio) long-read, 10X chromium library sequencing, accurate Illumina short read data and Hi-C library. We specifically described the diversity of transposable elements through a comparison with other oyster genomes and highlighted some specificities in non-coding RNA diversity.

## Materials and Methods

### Tissue sampling, RNA isolation, quality control and RNA-sequencing Tissues RNA for genome annotation

Samples of oyster tissues (gill, mantle, digestive gland, adductor muscle, gonads, hemocytes and palp) were sampled on 12 individuals and frozen in liquid nitrogen. Total RNA was extracted using TRIzol Reagent (Invitrogen, Life Technologies, USA) according to manufacturer’s instructions. The RNA quality was checked on agarose gel and quantified using a NanoPhotometer spectrophotometer (Thermo Fisher Scientific, Wilmington, DE). An equal amount of RNA of each tissue was pool to generate three libraries per tissue using 4 individuals per library. Libraries were constructed at the McGill Genome Center using NEBNext® Ultra™ Directional RNA Library Prep Kit for Illumina (New England Biolabs, Ipswich, MA, USA) following manufacturer’s recommendations and sequenced by Illumina HiSeq 2×150 cycles run (Illumina Inc., CA, USA).

### RNA extraction in *M. refringens* infected oyster

Fifty oysters (size: 5 to 8 cm) were collected at the banc du Roz (Bay of Brest, France). Hemolymph was retrieved in the adductor muscle using a syringe and hemocytes were collected by centrifugation and immediately frozen in liquid nitrogen. On the same individual, a piece of digestive gland was sampled for DNA extraction and parasite detection by PCR using two specific *M. refringens* primers (Leroux et al, 1999) and the remaining tissues were frozen in liquid nitrogen. RNA was then extracted from hemocytes and digestive gland of the six *M. refringens* positive individuals identified by PCR and on six negative individuals using TRIzol Reagent (Invitrogen). The RNA quality was checked on agarose gel and quantified using a NanoPhotometer spectrophotometer (Thermo Fisher Scientific). Two pools containing an equal amount of RNA from three samples for both *M. refringens* positive and negative samples were generated for RNAseq library construction and sequencing using the same protocol as described above and sequenced by Illumina HiSeq 2×150 cycles run (Illumina Inc., CA, USA).

### DNA library preparation and whole genome sequencing

#### DNA extraction

Genomic DNA used for PacBio, shot gun and 10X chromium libraries was extracted from a fresh adductor muscle dissected on a single specimen of *O. edulis* (size = 18 cm) collected in the Bay of Morlaix (Brittany, France). using a standard phenol-chloroform-isoamyl alcohol (PCI 25:24:1) protocol and ethanol precipitation. DNA quality was checked by electrophoresis on 1% agarose gels and DNA concentration was measured using a Qubit® dsDNA HS Assay Kit in Qubit® 2.0 Fluorometer (Life Technologies, CA, USA).

#### Shot-gun library

Shot-gun library was generated at the McGill Genome Center using the NxSeq® AmpFREE Low DNA Library Kit Library Preparation Kit (Lucigen Corp., WI, USA) according to the manufacturer’s recommendations and the library was ran on a HiSeq X for 2×150 cycles.

#### Pacbio Sequel libraries

The DNA library was prepared following the Pacific Biosciences 20 kb Template Preparation using 7.5 μg of high molecular weight genomic DNA using the SMRTbell Template Prep Kit 1.0 reagents (Pacific Biosciences, Menlo Park, CA, USA). The DNA library was size selected on a BluePippin system (Sage Science Inc., Beverly, MA, USA) using a cutoff range of 12 kb to 50 kb. The libraries were sequenced on a PacBio Sequel instrument at a loading concentration (on-plate) of 6 pM using the diffusion loading protocol, Sequel Sequencing Plate 2.1, SMRT cells 1M v2 and 10 hours movies.

#### 10X Chromium library

The gDNA was size selected on a BluePippin system (Sage Science Inc.) using a cutoff range of 40 kb to 80 kb. The 10x Chromium shotgun libraries were prepared following the Chromium Genome Reagent kits v2 User Guide RevB protocol, using the Chromium™ Genome Library & Gel Bead Kit v2, Chromium™ Genome Chip Kit and Chromium™ i7 Multiplex Kit (10X Genomics Inc., Pleasanton, CA, USA). The sequencing was performed on one lane of HiSeq X for 2×150 cycles.

#### Hi-C libraries

The Hi-C library construction protocol was adapted from Lieberman-Aiden et al., (2009) and Lazar-Stefanita (2017). Hi-C library was made from a piece of adductor muscle from another individual from the same population. Hi-C libraries were sequenced on a NextSeq 550 apparatus (2 × 75 bp, paired-end Illumina NextSeq). Contact maps were generated from reads using the hicstuff pipeline for processing generic 3C data, available at https://github.com/koszullab/hicstuff. The backend uses the bowtie2 (version 2.2.5) aligner run in paired-end mode (with the following options: --maxins 5 –very-sensitive-local). Alignments with mapping quality lower than 30 were discarded. The output was in the form of a sparsematrix where each fragment of every chromosome was given a unique identifier and every pair of fragments was given a contact count if it was nonzero. The assembled genome generated by instaGRAAL was polished to remove misassemblies (Baudry et al., 2020). Initial and final assembly metrics (Nx, GC distribution) were obtained using QUAST-LG. Misassemblies were quantified using QUAST-LG with the minimap2 aligner in the back-end. Ortholog completeness was computed with BUSCO (v3). Assembly completeness was also assessed with BUSCO. The evolution of genome metrics between cycles was obtained using instaGRAAL’s own implementation.

#### Gene prediction

All RNA-seq reads obtained from the different tissue transcriptomes were quality-trimmed using Trimmomatic (version 0.36) (Bolger et al., 2014) and mapped to the genome assembly using HISAT2 (Kim et al., 2019). The alignment information was processed to generate genome-guided transcriptome assembly using Trinity (ver. 2.8.4) (Grabherr et al., 2011) and in parallel, *de novo* transcriptome assembly was also generated by the Trinity software. The genome-guided and *de novo* transcriptome assemblies were processed by PASA pipeline (ver. 2.3.3) (Haas et al., 2003) to generate a training set for optimizing gene prediction parameters in AUGUSTUS (Stanke et al., 2006). RNA-seq reads were also aligned to the genome using STAR (ver. 2.7.1a) (Dobin et al., 2013) to produce hint files of exon and intron information. Then, *de novo* gene prediction was performed using AUGUSTUS (ver. 2.5.5). The completeness of the predicted gene models was assessed by Benchmarking Universal Single-Copy Orthologs (BUSCO) using metazoa_odb9 database (Simão et al., 2015). Gene annotation was made using Trinotate (Grabherr et al., 2011).

#### Noncoding RNA prediction

tRNA genes were predicted using tRNAscan-SE v1.3.1 (Lowe & Eddy, 1997) with eukaryote parameters, and rRNA with high conservation were predicted by aligning reads to the *Arabidopsis thaliana* template rRNA sequences using BLASTN (Altschul et al., 1997), with an e-value of 10^−5^. Additionally, INFERNAL (Nawrocki & Eddy, 2013) was used to predict miRNA and snRNA genes on the basis of the Rfam database (Griffiths-Jones, 2005).

### Repeat annotation

#### LTR-retrotransposons identification

We specifically investigated LTR-retrotranspons in five oyster genomes (*O. edulis, C. gigas* (GCA_902806645.1), *C. virginica* (GCA_002022765.4), *Pinctada martensii* (GCA_002216045.1) and *Saccostrea glomerata* (GCA_003671525.1)) using a detailed and precise pipeline previously customized for *M. mercenaria* (Farhat et al., 2022).

To annotate these LTR-retrotransposons, we first extracted and translated the RT/RNaseH domain from the sequences obtained with LTRHarvest using BLASTx (Camacho et al., 2009) (e-value less than 10^−5^) against an in-house database of RT/RNaseH (Thomas-Bulle et al., 2018) made from Gypsy Database (Llorens et al., 2011). Here, we kept the single sequences and/or a consensus sequence per previously defined-cluster and used them in phylogenetic approaches to determine the position of the elements in each clade. Cladistic analyses were performed on amino acid sequences corresponding to the RT/RNaseH domains of the newly characterized sequences and reference elements. Multiple alignments of these protein sequences were performed using MAFFT (Kato et al., 2018). After a manual curation of the alignments, phylogenetic analyses were conducted using Neighbor Joining and the pairwise deletion option of the MEGA5.2 software (Tamura et al., 2011). Using Topali2.3 (Milne et al., 2009), the best-fitted substitution model retained was the JTT model with a gamma distribution. Support for individual groups was evaluated with non-parametric bootstrapping using 100 replicates.

#### Repeated sequences annotation

Repeated sequences were annotated in the five oyster genomes by running RepeatMasker (http://repeatmasker.org) with default parameter and the script “Concatenate_sequences.py” (Thomas-Bulle et al., 2018). Two different libraries were used: (i) one with only inserts-cleaned consensus from each LTR-retrotransposon cluster detected by LTRHarvest to retrieve putatively missed copies (with corrupted LTRs or deleted); (ii) the second with all other consensus transposable elements obtained with RepeatModeler v2.0.1 (Flynn et al., 2020) using REPBASE, version 2017-01-27 (Jurka et al., 2005).

#### Differential Expression Analysis

RNAseq reads from *Martelia* libraries were checked for quality issues and adapter content with FastQC 0.11.7 and cleaning for sequencing adapters, trimming of low-quality bases (minimum mean quality score of 30), and filtering for length were performed with Trimmomatic 3.3 (Bolger et al., 2014). Reads were aligned to the genome assembly with Bowtie 2 (Langmead et al., 2009) to generate BAM files that were used to generate expression counts with the IdxStats software (Li et al., 2009). Raw read counts were imported into R 3.5.0 (R Core Team, 2017) for analysis with DESeq2 package (Love et al., 2014). The GO enrichment analysis was performed using DAVID software (Huang et al., 2009).

## Results

### Genome assembly and annotation

PacBio reads (45X raw coverage) and Illumina PE shotgun reads (104X raw coverage) were first assembled into contigs using Masurca hybrid assembler v3.2.8 (Zimin et al., 2017) assuming a 1 Gb genome size estimated by flux cytometry using SYBGREEN labelling (data not shown). Then, the 10X Chromium reads (67X raw coverage) were used by the ARCS v1.0.4 (https://github.com/bcgsc/arcs v1.0.4) together with LINKs pipeline v1.8.5 (https://github.com/bcgsc/LINKS, v1.8.5) to help scaffold and make the Masurca assembly more contiguous. A total of 5417 scaffolds were obtained for an estimated size of 1028 Mb with a N50 length of 0.94 Mb. HaploMerger2 software (https://github.com/mapleforest/HaploMerger2/releases/; Huang et al., 2017) was used to remove potential duplicated sequences and significantly improved the assembly to 2846 scaffolds (N50 of 1.62Mb). A chromosome-level assembly was generated using Hi-C data and InstaGRAAL software. The final assembly of the *Roscoff_O*.*edulis-V1* genome was 1018 Mb in size, with the chromosome-level scaffolds represented in 10 scaffolds corresponding to 887.2 Mb of sequence length and 553 unplaced scaffolds with a total N50 of 97.1 Mb for scaffold lengths. The 10 expected chromosomes are in accordance with the chromosomes previously evidenced in a karyotype analysis (Thiriot-Quiévreux & Ayraud, 1970) and linkage map (Lallias et al., 2007) (Supplementary Figure 1). The *Roscoff_O*.*edulis-V1* genome shows a GC content of 35.46% and the BUSCO analysis indicates that the gene models include 95.1% complete BUSCOs. A total of 35962 protein-coding genes were predicted from which 24302 genes were functionally annotated (e-value<0.001). The parameters of completeness and main genome features of the assembly are presented in Table 1 and are very close from those obtained in the *OE_Roslin_V1* assembly (Gundappa et al., 2022). Phylogenetic tree confirms that *Roscoff_O*.*edulis-V1* clusters with other ostreidae and has a closer relationship with the other two chinese ostrea species (Figure 1).

**Table 1.** Genome assembly statistics for *Roscoff_O*.*edulis-V1* genome.

| Metric | Value |  |
| --- | --- | --- |
| 10-chrom assembly size (bp) | 887 , 207 , 661 |  |
| Scaffold 1 (bp) | 117 , 936 , 894 | 4971 genes |
| Scaffold 2 (bp) | 111 , 702 , 126 | 4882 genes |
| Scaffold 3 (bp) | 97 , 982 , 138 | 4282 genes |
| Scaffold 4 (bp) | 97 , 658 , 951 | 4001 genes |
| Scaffold 5 (bp) | 97 , 091 , 390 | 4170 genes |
| Scaffold 6 (bp) | 95 , 771 , 753 | 3908 genes |
| Scaffold 7 (bp) | 90 , 095 , 513 | 3357 genes |
| Scaffold 8 (bp) | 74 , 472 , 875 | 2356 genes |
| Scaffold 9 (bp) | 53 , 076 , 851 | 2190 genes |
| Scaffold 10 (bp) | 51 , 419 , 170 | 1846 genes |
| GC content (%) | 35,46 |  |
| Complete BUSCOs* (C) | 931 (95.1%) |  |
| Complete and single-copy BUSCOs (S): | 911 (93.1%) |  |
| Complete and duplicated BUSCOs (D): | 20 (2.0%) |  |
| Protein coding genes number | 35963 |  |
| mean transcript length | 13916 |  |
| mean cds length | 1626 |  |
| mean exons per transcript | 7.2 |  |
| mean introns per transcript | 5.6 |  |

**Figure 1:**
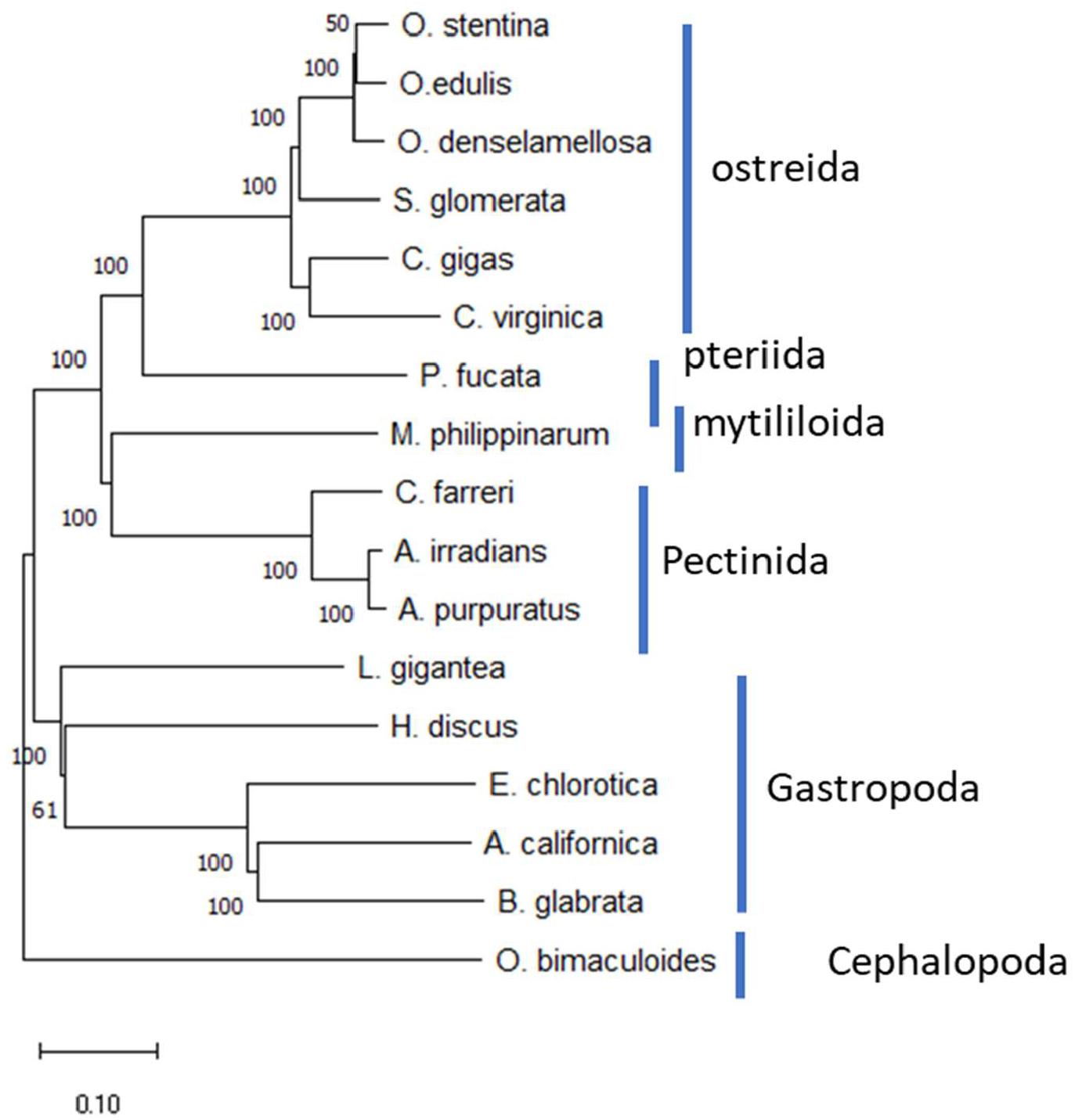
Phylogenetic position of the sequenced species. The phylogenetic tree was constructed based on a dataset from 114 single-copy orthologues generated from the aligment of the proteomes of *O. edulis, Argopecten irradians* (SRP174526), *A. purpuratus* PRJNA418203), *Aplysia californica* (GCA_000002075.2), *Biomphalaria glabrata* (GCF_000457365.1), *Crassostrea gigas* (GCA_902806645.1), *C. virginica* (GCA_002022765.4), *Chlamys farreri* (in MolluskDB), *Elysia chlorotica* (GCA_003991915.1), *Modiolus philippinarum* (GCA_002080025.1), *Pinctada fucata* (PRJNA283019), *Lottia gigantea* (PRJNA175706), *Octopus binaculoides* (GCF_001194135.1), *Saccostrea glomerata* (GCA_003671525.1), *Haliotis discus* (PRJNA317403), *Ostrea denselamellosa* and *O. stentina* (data specifically generated for this analysis) using Orthofinder version 2.4.1 (Emms & Kelly, 2019) with default parameters. MEGA version 11.7.8 software was used to construct the phylogenetic tree with bootstrap of 1,000 on a Neighbor Joining method.

### RNA prediction

Detection of miRNA using INFERNAL and Rfam database allowed the identification of 126 families of miRNAs (e-value < 10^−5^) in the *Roscoff_O*.*edulis-V1* genome. A large variation in the copy number is also detected among mi-RNA families and a total of 99 mi-RNA families contains more than one paralog. The mir-148 family is the most represented family and counts 219 copies followed by the mir-821 family with 106 copies and 23 families exhibiting more than 10 copies (Supplementary Table 1). The distribution of the 25 most represented mi-RNA families shows a relatively homogeneous distribution along the chromosomes with a higher frequency observed in chromosomes 5, 6, 7, 8 and 9. Most of the conserved mi-RNA families identified in other mollusc species (let-7, lin-4, miR-2, miR-29, miR-87, miR-184, miR-8, miR-10, miR-15, miR-17) are present in *Roscoff_O*.*edulis-V1* genome. We performed the same analysis on *C. virginica* and *M. mercenaria* genomes to identify enrichments in mi-RNA families in *O. edulis*. No mi-RNA family appears to be specific to *O. edulis* when comparing with *C. virginica* but *O. edulis* present a global higher number of copies for the most represented families (Table 2). In contrast, the diversity of mi-RNAs in *M. mercenaria* appears to be much lower and some families appear to be specific to oysters such as mir-148, mir-1027, mir-1803 and mir-355. Only mir-1253 is present in *M. mercenaria* and absent in oyster genomes (Table 2).

**Table 2:**
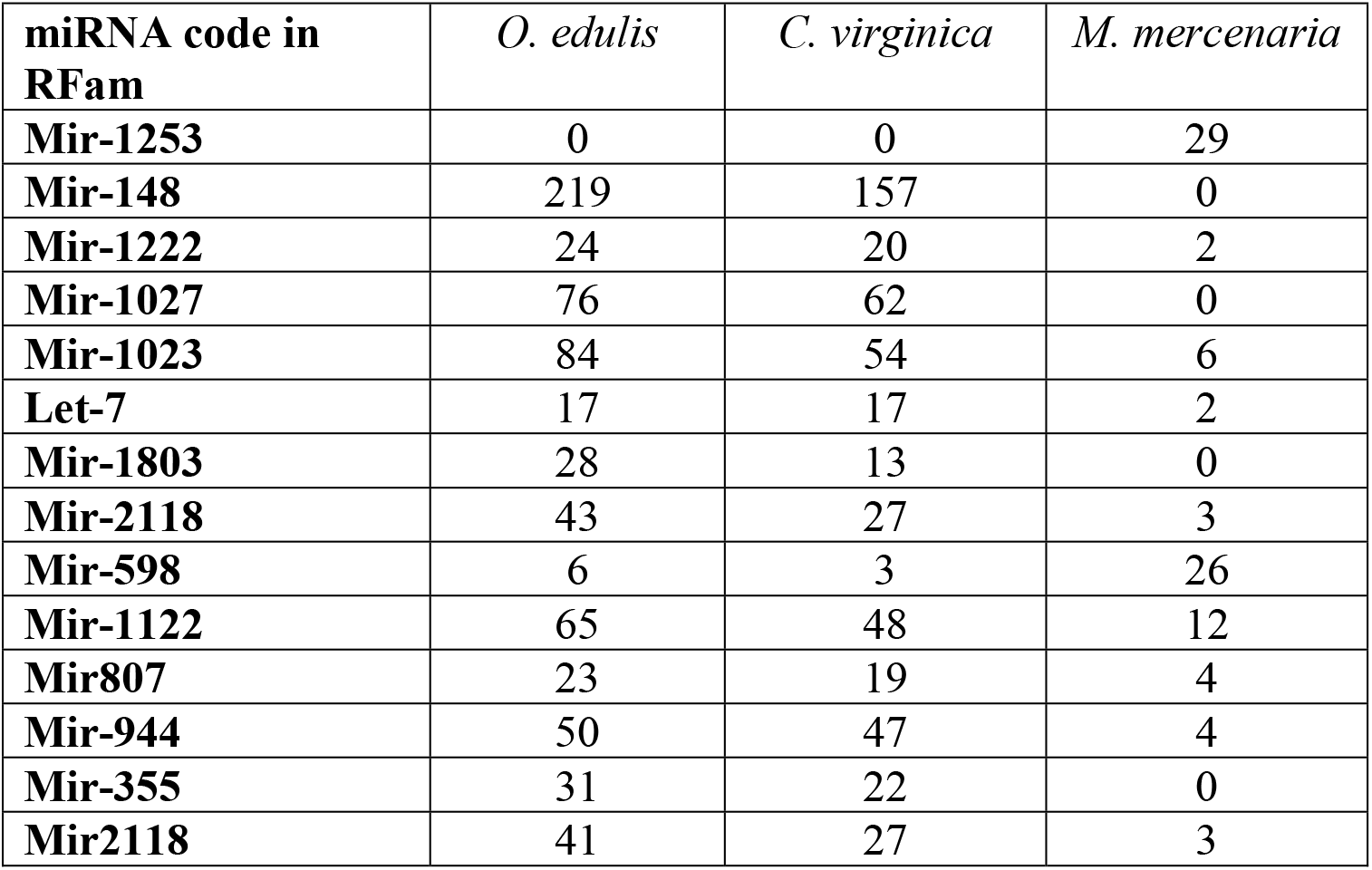
Numbers of copies for the 14 most represented miRNA in the genomes of *O. edulis, M. mercenaria* and *C. virginica (miRNA annotation according to RFam)*

A total of 1985 annotated tRNAs representing the 20 amino acids (Table 3), 3 copies of the 5.8S rRNA, 6 copies of the LSU-rRNA and 6 copies of SSU-rRNA but 54 copies of the 5S-rRNA have been annotated. Two tRNA families are over-represented with 1019 copies of the tRNA-Ser from which 907 correspond to anticodon AGA and 339 copies of the tRNA-Thr from which 282 copies correspond to AGU. tRNA-Ser gene distribution exhibits a homogenous repartition in the genome (Supplementary Figure 2). The pairwise alignment scores confirms that the tRNA-Ser motif are not included in a repeated element that could explain the high number of copies observed. The comparison of tRNA composition in *M. mercenaria* and *C. virginica* genomes using the same parameters showed that enrichment in tRNA-Ser was only detected in *O. edulis* but a higher content in tRNA-Thr (791 copies) and tRNA-Ala (465 copies) was detected in *M. mercenaria*. The density of tRNA is higher in *O. edulis* (2.275 tRNA/Mb) than in *M. mercenaria* (1.18 tRNA/Mb) or in *C. virginica* (0.575 tRNA/Mb). Comparison with available *C. gigas* tRNA composition show a similar pattern than with *C. virginica*.

**Table 3:** Numbers of tRNA and rRNA in the genomes of *O. edulis, M. mercenaria* and *C. virginica*

|  | *O. edulis* | *M. mercenaria* | *C. virginica* |
| --- | --- | --- | --- |
| tRNA_Ser | 1019 | 96 | 53 |
| tRNA_Phe | 42 | 25 | 11 |
| tRNA_Asp | 43 | 26 | 37 |
| tRNA_Glu | 31 | 20 | 56 |
| tRNA_iMet | 42 | 30 | 36 |
| tRNA_Tyr | 30 | 24 | 17 |
| tRNA_Leu | 44 | 51 | 60 |
| tRNA_Lys | 37 | 45 | 62 |
| tRNA_Gln | 39 | 48 | 21 |
| tRNA_Ile | 34 | 43 | 26 |
| tRNA_Cys | 20 | 26 | 10 |
| tRNA_Pro | 30 | 42 | 28 |
| tRNA_Arg | 49 | 74 | 95 |
| tRNA_Trp | 8 | 14 | 9 |
| tRNA-Sec | 1 | 2 | 1 |
| tRNA_Val | 28 | 56 | 34 |
| tRNA_His | 12 | 26 | 9 |
| tRNA_Thr | 339 | 791 | 153 |
| tRNA_Gly | 26 | 64 | 80 |
| tRNA_Met | 17 | 51 | 28 |
| tRNA_Asn | 21 | 72 | 21 |
| tRNA_Ala | 76 | 465 | 93 |
| **Total** | **1985** | **2091** | **940** |
| 5_8S_rRNA | 3 | 9 | 5 |
| 5S_rRNA | 54 | 18 | 99 |
| LSU_rRNA | 6 | 36 | 9 |
| SSU rRNA | 6 | 18 | 6 |
| **Total** | **69** | **81** | **119** |

**Table 4:** Copy number and genomic content of the repeated sequences in *O. edulis, C. gigas, C. virginica, P. martensii* and *S. glomerata* genomes generated following the annotation pipeline.

|  | *Ostrea edulis* |  |  | *Crassostrea gigas* |  |  | *Crassostrea virginica* |  |  | *Pinctada martensii* |  |  | *Saccostrea glomerata* |  |  |
| --- | --- | --- | --- | --- | --- | --- | --- | --- | --- | --- | --- | --- | --- | --- | --- |
|  | % genome | copies Nb | Nb of bp | % genome | copies Nb | Nb of bp | % genome | copies Nb | Nb of bp | % genome | copies Nb | Nb of bp | % genome | copies Nb | Nb of bp |
| **Retroelements** |  |  |  |  |  |  |  |  |  |  |  |  |  |  |  |
| **Penelope** | 4,46 | 57 561 | 38 234 354 | 0,49 | 4 421 | 3 174 681 | 0,11 | 764 | 740 415 | 5,84 | 90 093 | 57 830 428 | 3,01 | 39 094 | 23 757 514 |
| **LINEs:** | 3,05 | 31 673 | 26 178 491 | 2,83 | 18 815 | 18 348 219 | 1,79 | 12 916 | 12 273 678 | 0,94 | 10 497 | 9 359 382 | 1,53 | 12 860 | 12 064 399 |
| L2 | 0,43 | 4 915 | 3 721 701 | 0,04 | 310 | 239 915 | 0,03 | 237 | 210 627 | 0,09 | 1 076 | 925 824 | 0,28 | 1 812 | 2 188 757 |
| CR1 | 0,23 | 2 999 | 2 005 708 | 0,37 | 2 610 | 2 399 937 | 0,52 | 3 921 | 3 564 568 | 0,20 | 2 864 | 1 989 573 | 0,24 | 2 135 | 1 900 378 |
| RTE-X | 0,82 | 8 206 | 7 016 657 | 0,82 | 6 001 | 5 331 699 | 0,40 | 3 138 | 2 769 765 | 0,18 | 2 023 | 1 799 210 | 0,39 | 3 578 | 3 071 346 |
| I | 0,33 | 5 595 | 2 804 522 | 0,00 | 0 | 0 | 0,13 | 1 491 | 883 115 | 0,00 | 0 | 0 | 0,07 | 867 | 565 958 |
| L1 | 0,84 | 6 494 | 7 179 616 | 0,75 | 3 694 | 4 841 718 | 0,59 | 3 301 | 4 023 806 | 0,29 | 2 821 | 2 901 900 | 0,32 | 2 489 | 2 529 348 |
| RTE | 0,33 | 2 846 | 2 829 694 | 0,82 | 6 001 | 5 331 699 | 0,07 | 461 | 456 945 | 0,15 | 1 487 | 1 494 237 | 0,17 | 1 548 | 1 331 519 |
| Others | 0,07 | 618 | 620 593 | 0,03 | 199 | 203 251 | 0,05 | 367 | 364 852 | 0,03 | 226 | 248 638 | 0,06 | 431 | 477 093 |
| **SINEs** | 1,13 | 17 644 | 9 710 552 | 0,06 | 396 | 370 098 | 1,27 | 15 408 | 8 696 250 | 0,28 | 3 787 | 2 755 480 | 0,65 | 11 734 | 5 149 632 |
| **LTR elements:** | 1,61 | 9 362 | 14 030 935 | 2,51 | 10 438 | 16 258 283 | 1,63 | 8 440 | 11 191 940 | 1,00 | 6 838 | 9 867 404 | 1,94 | 11 497 | 15 278 733 |
| Gypsy | 1,33 | 8 012 | 11 588 206 | 2,07 | 9 159 | 13 397 133 | 1,41 | 7 519 | 9 650 060 | 0,88 | 6 273 | 8 757 301 | 1,70 | 10 292 | 13 403 278 |
| BEL/Pao | 0,26 | 1 237 | 2 229 921 | 0,42 | 1 231 | 2 734 597 | 0,21 | 795 | 1 410 415 | 0,07 | 503 | 706 175 | 0,22 | 1 144 | 1 745 192 |
| Copia | 0,02 | 79 | 189 857 | 0,02 | 48 | 126 553 | 0,01 | 45 | 100 528 | 0,00 | 0 | 0 | 0,02 | 61 | 130 263 |
| Others | 0,00 | 34 | 22 951 | 0,00 | 0 | 0 | 0,00 | 81 | 30 937 | 0,04 | 62 | 403 928 | 0,00 | 0 | 0 |
| **YR elements:** | 0,65 | 5 111 | 5 562 901 | 0,51 | 2 588 | 3 317 411 | 0,33 | 1 613 | 2 226 523 | 0,12 | 1 028 | 1 163 186 | 0,37 | 2 657 | 2 887 324 |
| Ngaro | 0,44 | 3 150 | 3 810 113 | 0,3442 | 1 768 | 2 230 023 | 0,19 | 982 | 1 287 955 | 0,02 | 196 | 212 583 | 0,27 | 1 810 | 2 115 549 |
| DIRS | 0,20 | 1 961 | 1 752 788 | 0,1678 | 820 | 1 087 388 | 0,14 | 631 | 938 568 | 0,10 | 832 | 950 603 | 0,10 | 847 | 771 775 |
| **DNA transposons** | 7,08 | 83 982 | 60 748 905 | 7,32 | 74 071 | 47 420 818 | 5,34 | 46 434 | 36 532 449 | 3,56 | 57 625 | 35 264 480 | 5,88 | 69 471 | 46 338 311 |
| Maverick | 0,28 | 751 | 2 438 022 | 0,22 | 764 | 1 393 542 | 0,38 | 1 133 | 2 580 104 | 0,19 | 991 | 1 897 408 | 0,08 | 701 | 600 613 |
| TcMar/Pogo | 1,38 | 14 389 | 11 848 554 | 1,19 | 12 709 | 7 710 105 | 0,49 | 4 683 | 3 333 461 | 0,48 | 7 725 | 4 741 920 | 1,38 | 13 911 | 10 870 057 |
| hobo/Ac/Tam | 0,95 | 12 619 | 8 173 678 | 0,39 | 3 002 | 2 520 162 | 0,46 | 4 165 | 3 166 922 | 0,80 | 12 909 | 7 915 023 | 0,49 | 5 663 | 3 866 272 |
| Zator | 0,38 | 4 849 | 3 295 890 | 0,15 | 1 481 | 960 144 | 0,04 | 425 | 293 547 | 0,15 | 3 014 | 1 487 162 | 0,14 | 1 896 | 1 064 820 |
| Crypton | 1,37 | 20 899 | 11 730 201 | 1,78 | 22 914 | 11 503 648 | 1,09 | 12 912 | 7 489 797 | 0,75 | 15 751 | 7 448 289 | 1,32 | 19 279 | 10 442 231 |
| Academ | 0,25 | 2 649 | 2 153 984 | 0,03 | 167 | 213 307 | 0,13 | 651 | 894 907 | 0,08 | 791 | 743 310 | 0,03 | 229 | 274 905 |
| EnSpm | 0,09 | 705 | 809 979 | 0,32 | 1 788 | 2 080 891 | 0,47 | 3 155 | 3 220 923 | 0,16 | 2 902 | 1 579 436 | 0,31 | 2 877 | 2 450 110 |
| MULE/MuDR/IS905 | 0,13 | 1 341 | 1 130 079 | 0,04 | 328 | 288 206 | 0,10 | 806 | 672 632 | 0,07 | 875 | 678 893 | 0,08 | 701 | 600 613 |
| PHIS | 1,22 | 12 327 | 10 474 972 | 1,56 | 14 069 | 10 077 041 | 1,14 | 9 623 | 7 838 419 | 0,66 | 10 261 | 6 499 076 | 1,09 | 13 198 | 8 619 218 |
| Kolobok | 0,58 | 8 162 | 5 001 169 | 1,24 | 14 339 | 8 041 774 | 0,61 | 6 107 | 4 177 778 | 0,03 | 385 | 334 047 | 0,65 | 7 877 | 5 160 850 |
| PiggyBac | 0,11 | 1 710 | 965 878 | 0,15 | 823 | 962 318 | 0,13 | 485 | 901 089 | 0,00 | 0 | 0 | 0,09 | 1 064 | 748 137 |
| Sola | 0,16 | 1 741 | 1 401 559 | 0,15 | 896 | 988 546 | 0,13 | 855 | 871 136 | 0,11 | 990 | 1 119 636 | 0,09 | 673 | 703 386 |
| others | 0,15 | 1 840 | 1 324 940 | 0,11 | 791 | 681 134 | 0,16 | 1 434 | 1 091 734 | 0,08 | 1 031 | 820 280 | 0,12 | 1 402 | 937 099 |
| **Rolling-circles** | 9,88 | 110 094 | 84 753 110 | 12,47 | 79 436 | 80 781 395 | 9,48 | 72 591 | 64 896 885 | 3,83 | 54 233 | 37 948 915 | 9,88 | 110 094 | 84 753 110 |
| **Unclassified** | 10,20 | 131 922 | 87 508 527 | 3,90 | 31 005 | 25 285 376 | 8,74 | 82 096 | 59 864 032 | 0,09 | 1 296 | 867 014 | 11,24 | 131 498 | 88 620 649 |

### Transposable elements in *O. edulis* and other oyster genomes

Repeated elements were annotated in the five assemblies using the same pipeline method for proper comparison (Table 5) showing a similar content of various repeated elements, except for *P. martensii* for which TEs represent less than 15% of the genome. Amon the five species, *O. edulis* presents the highest coverage in TEs (38.05%). Although each element type is variably represented among species (e.g., Penelope elements are rare in both Crassostrea, as are SINEs in *C. gigas* and *P. martensii*), *O. edulis* almost always has the highest (LINEs, YR elements) or second-highest (Penelope, SINEs, DNA transposons, rolling circles) proportions of TEs for each element type. Only LTR-retrotransposons seem to be a little less frequent (1.61%) than in other species.

**Table 5:** Genomic proportions of the different clades of LTR-retrotransposons detected in *O. edulis, C. gigas, C. virginica, P. martensii* and *S. glomerata* genomes. The number of sub-families of each clade is given in brackets. “Unclassified” = elements not linked to a clade; “-” = no element detected; “s” = only single sequences detected:

|  | *O. edulis* | *C. gigas* | *C. virginica* | *P. martensii* | *S. glomerata* |
| --- | --- | --- | --- | --- | --- |
|  | 872 374 424 bp | 647 887 097 bp | 684 723 884 bp | 990 984 031 bp | 788 100 799 bp |
| **GalEa** | **0.022 (4)** | **0.020 (4)** | **0.015 (2)** | **s** | **0.017 (4)** |
| Flow | s | 0.015 (1) | - | 0.003 (1) | s |
| Suzu | 0.005 (1) | 0.015 (2) | 0.006 (1) | s | 0.006 (1) |
| Tas | 0.027 (4) | 0.019 (2) | 0.041 (3) | - | - |
| Sinbad | 0.034 (5) | 0.036 (3) | 0.027 (2) | 0.004 (2) | 0.015 (2) |
| Sparrow | 0.187 (26) | 0.300 (33) | 0.127 (16) | 0.055 (9) | 0.161 (25) |
| Surcouf | s | 0.021 (2) | 0.005 (1) | 0.004 (1) | 0.011 (2) |
| Unclassified | 0.002 (1) | 0.017 (2) | - | 0.005 (1) | 0.029 (6) |
| **Pao/BEL** | **0,256** | **0,422** | **0,206** | **0,071** | **0,221** |
| AB-clade | 0.034 (6) | 0.039 (7) | 0.025 (3) | s | 0.034 (8) |
| C-clade | 0.142 (28) | 0.117 (13) | 0.138 (18) | 0.053 (16) | 0.133 (23) |
| MolGy1 | 0.347 (23) | 0.376 (32) | 0.290 (25) | 0.075 (6) | 0.319 (26) |
| MolGy2 | 0.584 (47) | 0.631 (51) | 0.450 (29) | 0.208 (21) | 0.672 (52) |
| MolGy3 | 0.062 (11) | 0.039 (5) | 0.039 (6) | 0.024 (6) | 0.053 (10) |
| MolGy4 | 0.014 (1) | 0.250 (7) | s | 0.213 (4) | 0.044 (2) |
| MolGy5 | 0.050 (2) | 0.005 (1) | 0.099 (2) | 0.220 (8) | 0.121 (3) |
| MolGy6 | - | 0.216 (11) | 0.079 (6) | 0.018 (2) | 0.070 (4) |
| MolGy10 | 0.006 (1) | 0.018 (1) | 0.011 (1) | - | 0.013 (2) |
| MolGy17 | 0.0182 (2) | s | s | 0.007 (1) | 0.011 (2) |
| MolGy18 | 0.005 (1) | 0.006 (1) | - | 0.003 (1) | 0.010 (1) |
| MolAn | 0.026 (2) | 0.262 (16) | 0.223 (10) | s | 0.132 (14) |
| Unclassified | 0.041 (7) | 0.109 (8) | 0.056 (9) | 0.064 (12) | 0.089 (11) |
| **Gypsy** | **1,328** | **2,068** | **1,409** | **0,884** | **1,701** |
| **LTR-retrotransposons** | **1,606** | **2,509** | **1,630** | **0,955** | **1,939** |

In order to characterize LTR-retrotransposons in oyster genomes, LTRharvest was used and its output was integrated in phylogenetic analyses. For this purpose, two types of sequences were used: (i) either a consensus devoid of possible insertions in the case of a cluster of elements previously defined by Uclust, (ii) or isolated (single) sequences that did not cluster with any other elements. A total of 797 sub-families were detected by LTRharvest in the five genomes, including 14 Copia, 155 BEL/Pao and 628 Gypsy (Supplementary Table 2). The relative abundance of the three superfamilies in *O. edulis* (76% for Gypsy elements, 22% for BEL/Pao elements and 2% for Copia elements) is quite similar between oyster species; except for Copia elements in *P. martensii* only represented by one single copy (Table 5). Phylogenetic tree of Copia elements revealed that all elements only belong to the GalEa clades (Figure 2).

**Figure 2:**
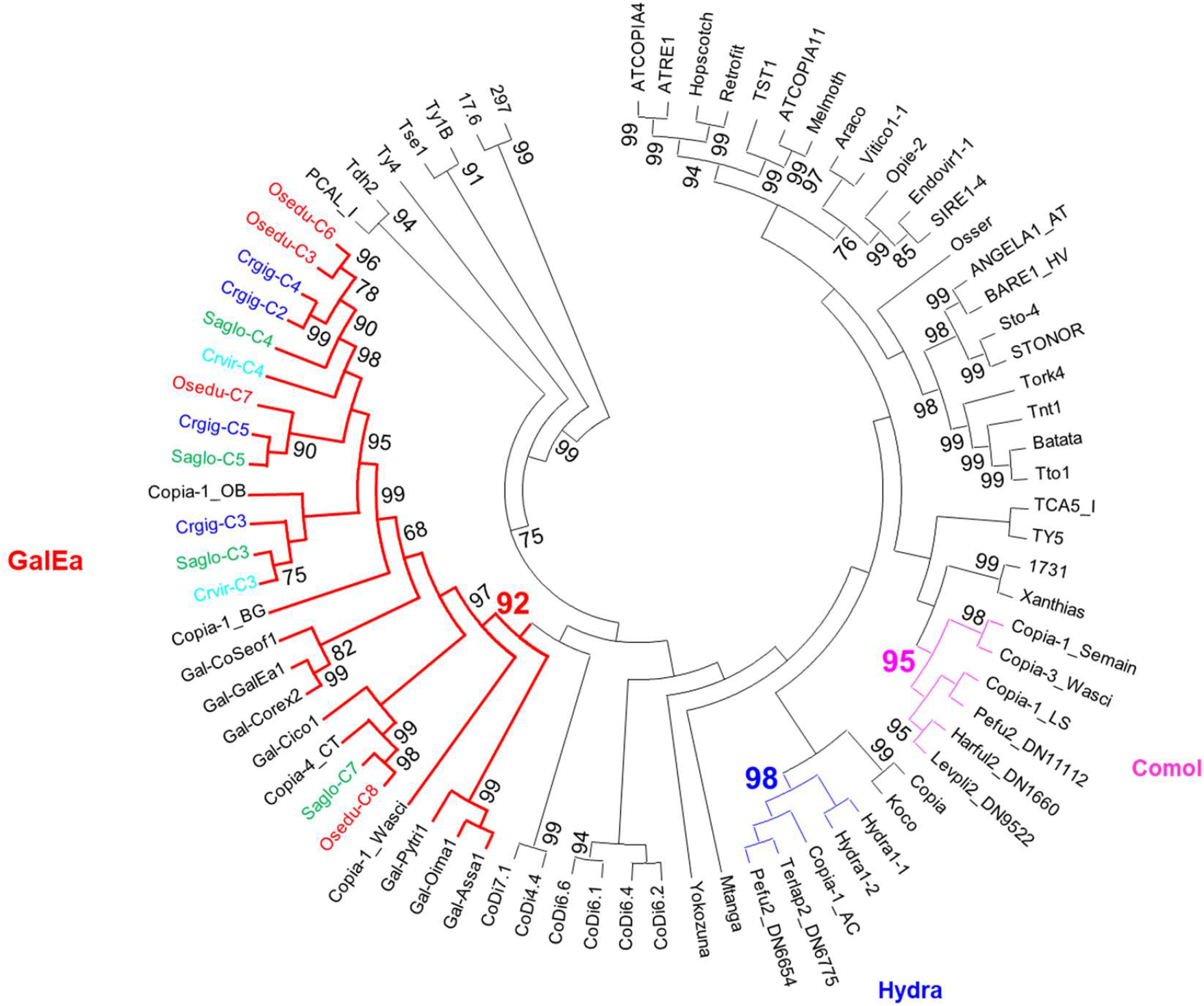
Phylogenetic relationships of Copia retrotransposons. The tree is based on Neighbor-Joining analysis of RT/RNaseH domain amino acid sequences. The Copia sub-families from oysters are indicated in color (*Crassostrea gigas* in dark blue, *Crassostrea virginica* in light blue, *Ostrea edulis* in red, *Pinctada martensii* in purple and *Saccostrea glomerata* in green) as are the clades known to possess elements of mollusks. Node statistical support values (>70 %) come from non-parametric bootstrapping using 100 replicates.

*O. edulis* genome did not contain elements of the BEL, Pao, and Dan clades (Supplementary Figure 3) but contains TEs in the six other clades in the BEL/Pao superfamily. Most of its elements belong to the Sailor lineage, mostly to the Sparrow, Sinbad and Tas clades and only one sub-family in the Suzu clade and one element in the Flow clade are present. This distribution is globally observed in the other oysters, with a large Sparrow clade. Only *O. edulis* and *C. gigas* present all the six clades; with in particular the absence of element of the Tas clade in both *P. martensii* and *S. glomerata*.

Gypsy superfamily tree reveals 12 clades in oysters (Figure 3, Supplementary Figure 4). Ten correspond to the MolGy clades previously defined from mollusk elements (except for clades MolGy11 and MolGy13) and to the clades A-B and C, also known to have Gypsy elements of mollusks. Two new putative clades can be further identified with oyster elements: MolGy17, observed in all 5 species and MolGy18 which is only absent in *C. virginica. O. edulis* do not present any element of the clade MolGy6, although it is quite well represented in the 4 other species; and it also has very few elements of the MolAn clade (Table 5). Three clades (Molgy1, Molgy2 and the C-clade) are well recovered, represent about 1.26 % of the genome and largely dominate with three quarters of the sub-families as observed in other oysters.

**Figure 3:**
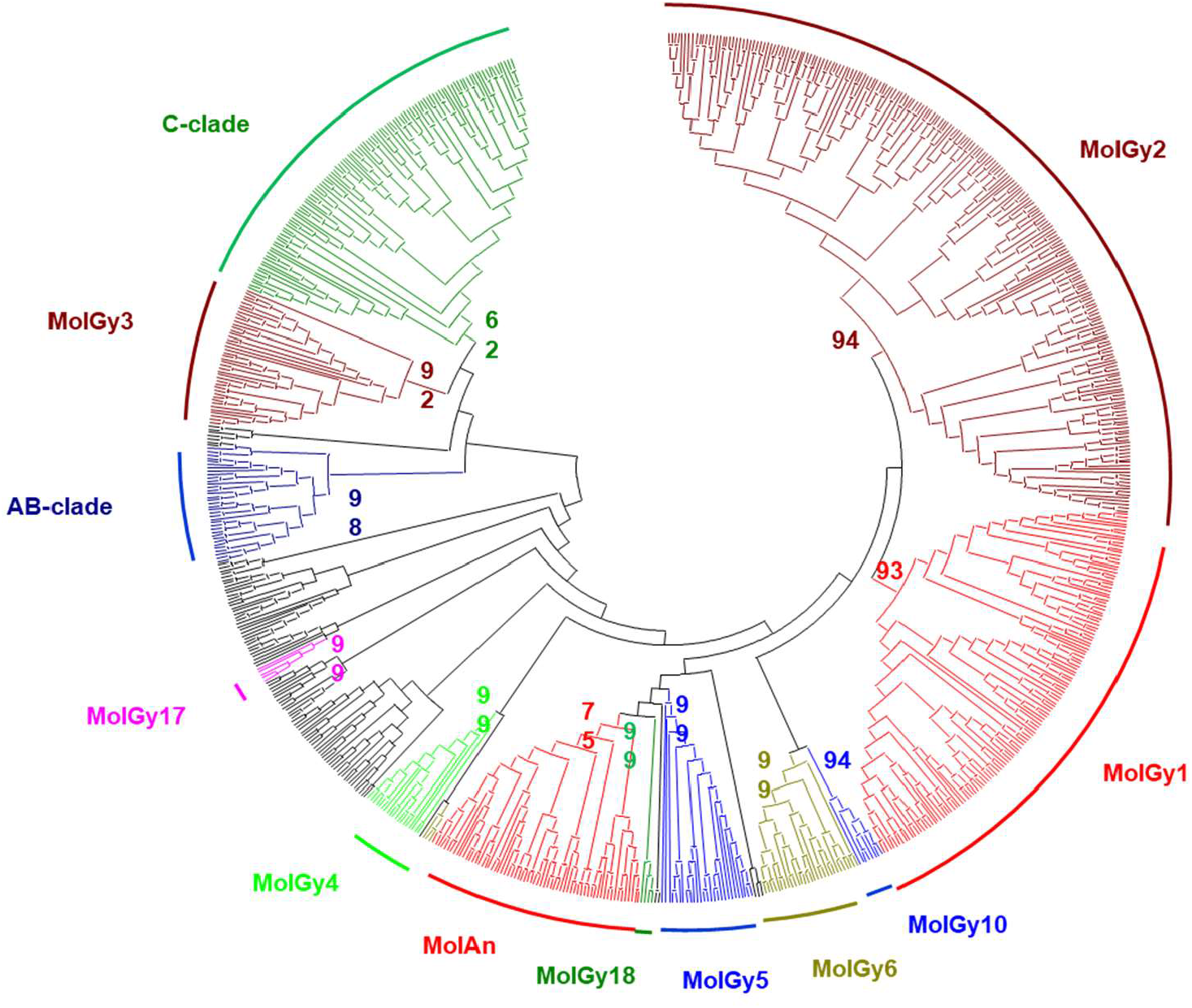
Phylogenetic relationships of Gypsy retrotransposons. The tree is based on Neighbor-Joining analysis of RT/RNaseH domain amino acid sequences. The clades that have elements of oysters are indicated in color. Node statistical support values (>70 %) come from non-parametric bootstrapping using 100 replicates.

### Differential Expression Analysis in *Marteilia infected oysters*

In the hemocytes, 143 DEGs are down-regulated in *Martelia* infected oyster (Figure 4). GOterm enrichment analysis show that 4 GOterms into the Biological Process category (GO:0007155 cell adhesion, GO:0032715 negative regulation of interleukin-6 production, GO:0032088 negative regulation of NF-kappaB transcription factor and GO:0070373 negative regulation of ERK1 and ERK2 cascade), 5 GOterms in the Cellular Component category (GO:0005581 collagen trimer, GO:0005576 extracellular region, GO:0005615 extracellular space, GO:0031012 extracellular matrix and GO:0030054 cell junction) and 3 GOterms in the Molecular Function category (GO:0005102 receptor binding, GO:0030020 extracellular matrix structural constituent conferring tensile strength and GO:0005518 collagen binding) are significantly over-represented (Supplementary Table 3). 78 DEGs are up-regulated and 4 GOterms into the Biological Process category (GO:0006259 DNA metabolic process, GO:0090305 nucleic acid phosphodiester bond hydrolysis, GO:0006302 double-strand break repair, GO:0007155 cell adhesion), and 2 GOterms in the Molecular Function category (GO:0003682 Chromatin Binding and GO:0003677 DNA binding) are enriched (Supplementary Table 4). Among the most down-regulated regulated genes, we identified IAPs, mannose receptor C-type, serine protease inhibitor, one SOD and several genes encoding C1q complements which are all involved in immune response. Among up-regulated genes, we identified several proteins involved in apoptosis such as ced-1 that are up-regulated and other genes involved in cell adhesion such as a sushi, von Willebrand factor type A, an EGF and the pentraxin domain containing protein 1 and several TRIM proteins.

**Figure 4:**
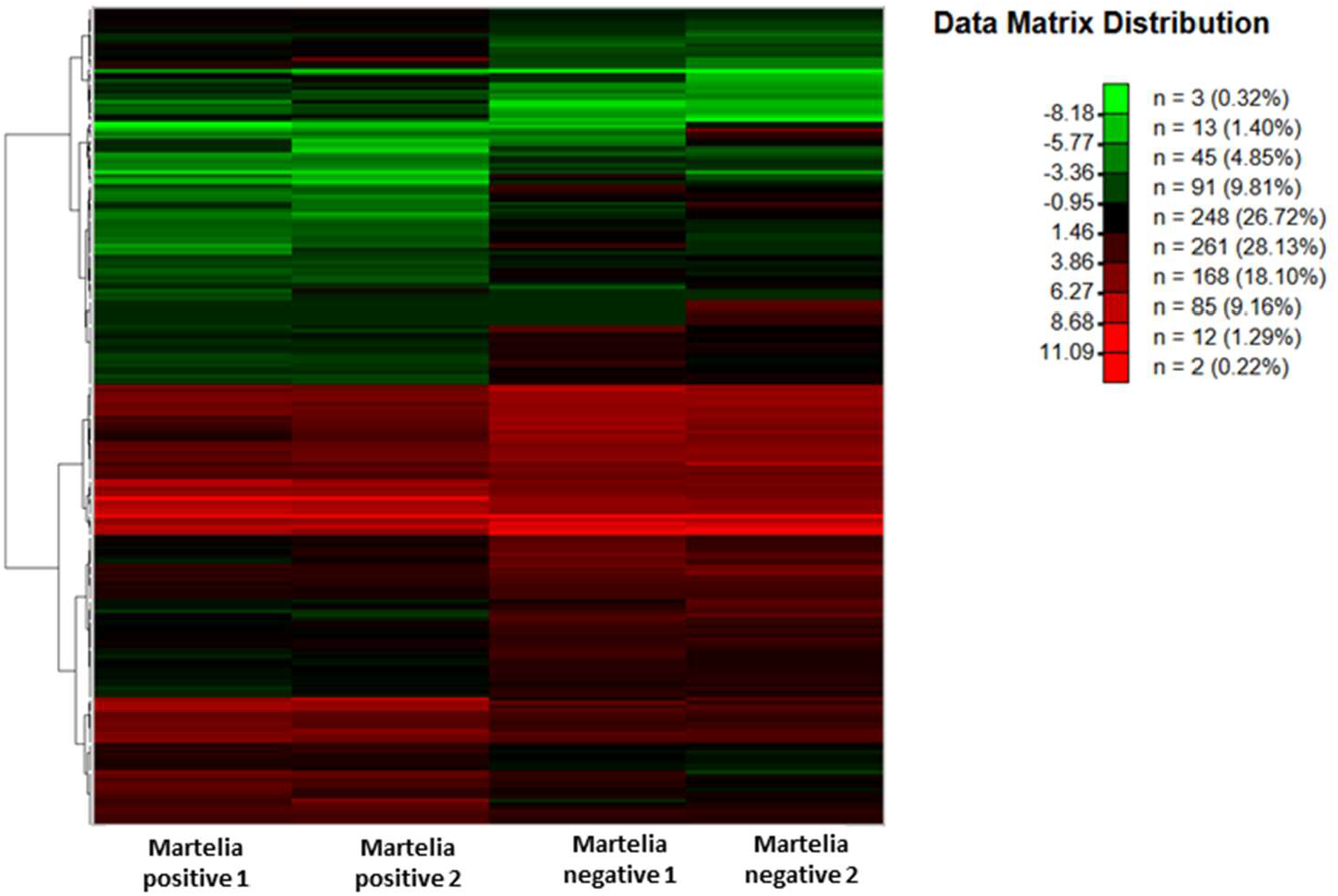
Hierarchical clustering of DEGs in hemocytes of Martelia infected and non infected *O. edulis* individuals (the data matrix distribution represents the log2 intensity values)

In the digestive gland, a total of 926 DEGs have been identified as over-expressed in *Martelia* infected oysters (Figure 5) and represent 29 terms into the Biological Process category, 28 terms in the Cellular Component category and 12 terms in the Molecular Function category (Supplementary Table 5). 1640 DEGs have been identified as down-regulated in *Martelia* infected oysters representing 68 GOterms into the Biological Process category, 22 GOterms in the Cellular Component category and 33 GOterms in the Molecular Function category (Supplementary Table 6). Among the genes showing the strongest up-regulations in *M. refringens*-infected individuals (log2 fold > 4), we identified several genes known to be involved in immune response processes or cancer-like pathologies (prominin, DMBT1, 2-proprotein convertase subtilisin/kexin), cell adhesion (contactin, laminin). Several genes involved in DNA, RNA or histone methylation process are also shown to be up-regulated (log2 fold >2.5). At the opposite, three genes encoding for Cadherin 12 that are involved in cell adhesion homeostasis, one gene involved in the protection of germline (Mage protein), hemocyte aggregation (hemocytin), RNA editing (ADAR) are strongly down-regulated. Several genes involved in the positive regulation of transcription from RNA polymerase II promoter and a total of 14 genes encoding G-protein-coupled receptors are also down-regulated

**Figure 5:**
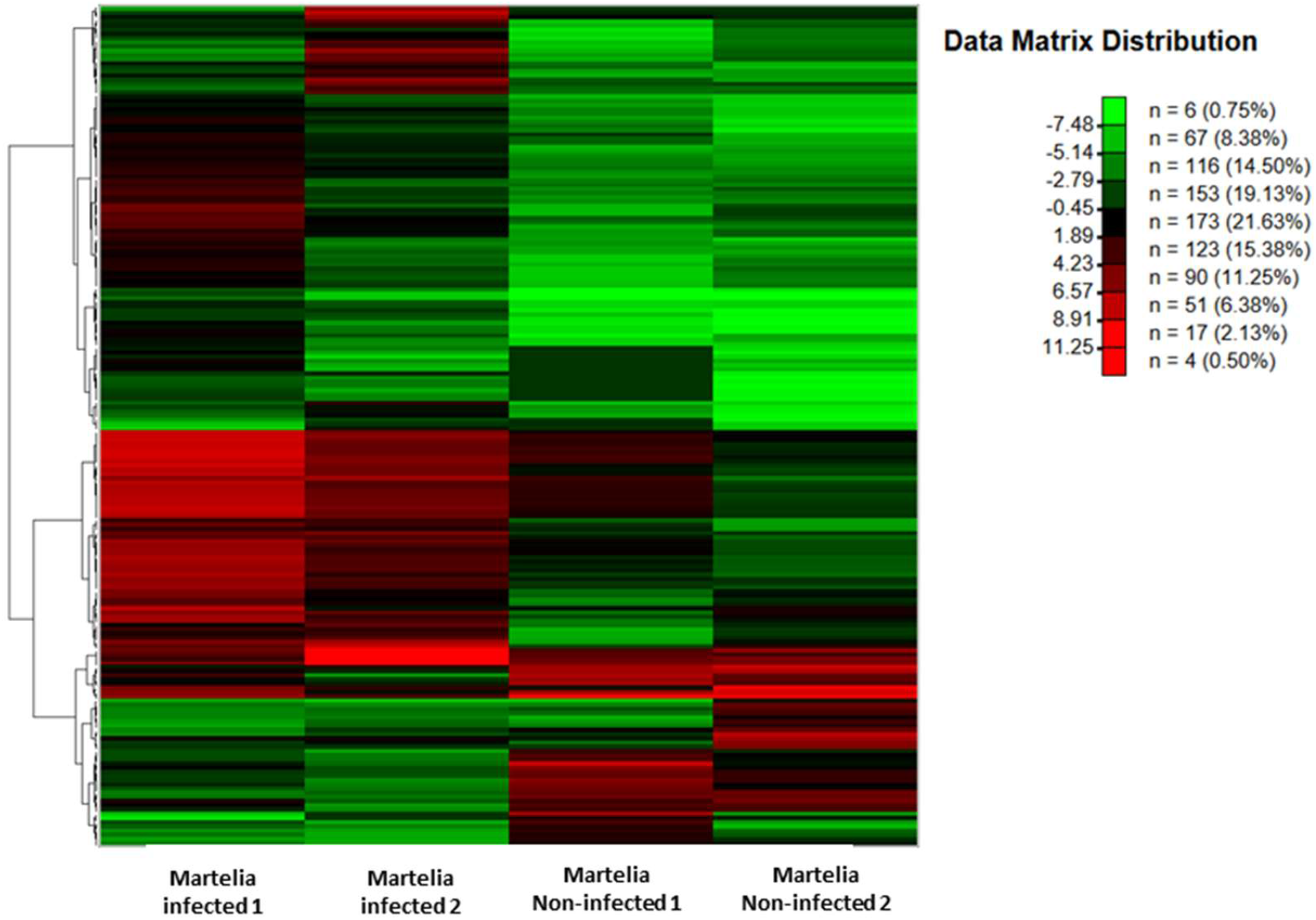
Hierarchical clustering of DEGs in digestive glands of Martelia infected and non infected *O. edulis* individuals (the data matrix distribution represents the log2 intensity values)

## DISCUSSION

The use of the combined long reads, shot gun and 10X sequencing approaches allowed us to a obtain a high quality reference genome assembled at a chromosomal level for the flat oyster O. eduliswas made available for subsequent genomic analysis. With a size of 887 Mb distributed along the 10 predicted chromosomes and good assembly statistics, the *Roscoff_O*.*edulis-V1* assembly genome ranks among other sequenced genomes of bivalve mollusks (Li et al., 2021; Peñaloza et al., 2021; Farhat et al., 2022). The *Roscoff_O*.*edulis-V1* genome size is larger that of the Crassostreidae (around 600Mb) but the number of genes predicted is close to that obtained in oyster species such *C. gigas* (31371 proteins), *C. virginica* (34 587 proteins) or *P. fucata* (31477 proteins).

### Analysis of tRNA and miRNA reveales specificity in *Ostrea* genome

Transfer RNAs play an essential role in cellular life and in many metabolic processes (Francklyn & Minajigi, 2010). In many species, the number of genes encoding each tRNA generally correlates with amino acid frequency (Duret, 2000) with some exceptions such as in Arabidopsis thaliana, where a higher number of genes encoding tRNAs-Tyr, Ser and Pro are detected (Michaud et al., 2011). Counting of serine residues in the *O. edulis* proteome does not reveal an excess of this residue over other amino acids, failing to explain the high tRNA-Ser copy number. The counting of serine amino acids in the proteomes of three other mollusks (*C. gigas, C. virginica* and *M. mercenaria*) does not show any significant difference (from 7.9 to 8.28% depending on the species). The same result was obtained when considering the proportion of threonine residues in proteome despite a high number of tRNA-Thr genes in *O. edulis*. The number of tRNA genes identified in the three species we analyzed is consistent with what is found in other mollusk species such as *Scapharca broughtonii* with 1541 tRNA genes (Bai et al., 2019) or *C. gigas* with 641 tRNA genes (Peñaloza et al., 2021). In eukaryotes, no general trend in the number of tRNA genes is evidenced and the strong variations observed could reflect differences in the evolutionary history of lineages. While the study of tRNA genomic organization and structural evolution remains partial (Marck & Grosjean, 2002; Goodenbour & Pan, 2006), tRNA genes have been shown to play a role in genome organization by acting as barriers to DNA replication fork progression (McFarlane & Whitehall, 2009) or by contributing to chromosomal instability (Admire et al., 2006). No clear relation between tDNA copy number and their organisation in clusters in genomes is neither evidenced and most tDNA clusters are small, containing only a few colocalized tRNA genes with the exception of Nematostella and some fishes species (Bermudez-Santana et al., 2010). Despite a high number of the Ser-tRNA-Ser and tRNA-Thr which also present a high disequilibrium in the proportion of isoacceptors in the *Roscoff_O*.*edulis-V1* genome, no specific clustering of those tRNA genes is observed. Variations in codon usage might influence the variation of tDNA copy numbers (Rocha, 2004) but no strong desequilibrium for Ser resides has been evidenced in *O. edulis* (Gerdol et al., 2015). Differences in the number of rDNA loci between individuals of the same species (ie *M. galloprovincialis*), as well as in the location of the rDNA loci between different cells from the same individual has been evidenced (Insua and Mendez, 1998). Additional analyses have to be address to better understand if and how the enrichment observed for the tRNA-Ser and tRNA-Thr has a functional role in *O. edulis* physiology because this specific pattern is not observed in other oysters genomes and no difference in amino acid composition has been detected between proteomes.

Despite that mollusks represent the second largest phylum in Metazoa, only a very limited number of studies concerning the identification and the role of miRNA has been addressed despite their essential role in cellular development, proliferation, apoptosis, oncogenesis, differentiation, and disease (Bueno and Malumbres, 2011). If more than 48800 miRNA are available in all species (www.mirbase.org), only 245 families have been identified in molluscs mainly due to the low number of genomes available, a lack of specific miRNA annotation and a very limited number of miRNA transcriptomic analysis. The number of miRNA families (128) identified in *O. edulis* are in agreement with what is observed in *C. gigas* (120 families), *C. virginica* (121 families) or *P. fucata* (126 families) and many of the mi-RNA classically found in molluscs are present. The comparison of the miRNAs copy number shows differences within a species between the different approaches used for their annotation limiting the ability to make reliable comparisons. In *C. gigas*, 8 copies of miRNA-184 are identified by Huang et al., (2021) versus 44 by Rosani et al., (2021). More, differences are evidenced when comparing predicted miRNA and miRNA identified in small-RNAseq sequencing which do not confirm that the predicted miRNAs are actually expressed. In the *Roscoff_O*.*edulis-V1*, we identified several miRNA families that exhibit a high numbers of copies distributed along the 10 chromosomes without specific clustering. The most represented family is the miRNA-148 with 219 copies and is known to act as a negative regulator of MyD88-dependent NF-κB signaling in the teleost fish (Chu et al., 2017) and which is inhibited in the herbivorous carp *Ctenopharyngodon idella*, in response to bacterial infection (Fang et al., 2020). The Mir-841 gene with 106 copies in *O. edulis genome* is known to be involved in the plants in response to metabolic stress (Nischal et al., 2014; Lu et al., 2014) or virus (Moyo et al., 2017). The involvement of several miRNAs in response to the herpes virus OsHV-1 have been demonstrated in *C. gigas* (Rosani et al., 2020), *Chlamys farreri* (Chen et al., 2014). In *O. edulis*, miRNA involved in immune system regulation (mi-RNA1, miRNA10, miRNA8, mi-RNA33, have been shown to be regulated in response to *B. ostreae* (Martin-Gomez et al., 2014). A specific analysis has to be conducted on *O. edulis* by coupling mi-RNA prediction and miRNA transcriptomic profiling.

### Transposable elements

Comparison of the TEs composition in the five oyster species analyzed shows a relative homogeneity of their content. The percentages of genome coverage are very close for most types of elements (LINEs, SINEs, DNA transposons, YR elements) and the only remarkable difference in *O. edulis* is the over-representation of Penelope elements (4.46% vs. 0.5% in other species), and a global under-representation of LTR-retrotransposons (1.61% in *O. edulis* vs. 4.37%). With the exception of the MolGy6 clade, *O. edulis* possess all the clades detected in oysters and 17 of the 23 clades described in mollusks are present in oysters. A targeted study of Gypsy elements within 20 bivalve genomes revealed 20 branches within the Gypsy clade C (Farhat et al., 2022), and 18 are found in oyster genomes. Analysis of the 5 oyster genomes has further defined two new clades, MolGy17 and MolGy 18 that appear to be specific to oysters. Specific elements such as the Steamer LTR-retrotransposon family which is associated to neoplasia in other bivalve species (Metzger et al., 2018) have not be detected in *O. edulis* genome while it is present in other oysters like *C. gigas, C. virginica* and *S. glomerata* (Farhat et al., 2022). Contrary to *C. gigas*, no specific enrichment in the rolling-circle transposable elements Helitrons is detected in *O. edulis* as well as the steamer element which is present in Crassostrea species. However, the high percentage of unclassified TEs in both *O. edulis* and *S. glomerate* genomes suggests that flat oysters could contain new families of ETs not yet described. Transposable elements (TEs) are known to have a large impact on genome structure and stability, and are therefore considered to play an important role in many evolutionary and ecological processes such as biodiversity generation, stress response, adaptation, speciation and colonization mechanisms evolution in eukaryotes (Kazasian, 2004; Biemont & Vieira, 2006; Finnegan, 2012). Environmental variations can promote genome plasticity through transcriptional activation and TEs mobilization, often in response to specific stimuli such as biotic stresses (pathogen) or abiotic environmental changes (temperature, heavy metals, UV) (Bennetzen, 2000; Capy et al., 2000; Melayah et al., 2001). This variability transmitted to the offspring would also favor a better adaptation of the organisms to environmental changes (long-term effect detectable in the genome). Extensive analysis on TEs regulation in response to biotic and abiotic environmental parameters has to be conducted in better understand the role of TE in the flat oyster biology.

### Transcriptomic regulation in response to *Martelia*

Studies of the responses of Ostrea to the presence of Martelia have not yet been carried out, unlike B. ostreae, despite the strong impact of Martelia in terms of mortality on the Ostrea beds and more particularly on the foreshore. This first study carried out on individuals naturally infected by *M. refringens*, although preliminary, allowed to highlight some metabolic pathways classicaly involved in parsite interaction but also new pathways that could be more specifically associated to the response to a eukaryote parasite. Indeed, in hemocytes, the main tissue involved in immune response, the down-regulation of genes such as IAPs, mannose receptor C-type, serine protease inhibitor, and C1q complement that participate to the molecular response to various parasites (Li et al., 2019; Chan et al., 2021). The down-regulation of those genes could indicate a lower capacity of immune response by the oyster. C1q domain containing proteins have been largely characterized in mollusk genomes, are characterized by a large diversity gene number across species (Mun et al., 2017; Gerdol et al., 2019, Peng et al., 2020; Farhat et al., 2022) and have been shown to be overrepresented in *O. edulis* genome (Gundappa et al., 2022the over-expression of genes involved in apoptosis is coherent with previous results obtained on *O. edulis* in response to *B. ostreae* (Gervais et al., 2019) or in *C*.*virginica* in response to *Perkinsus marinus* (Lau et al., 2018)

The high number of differentially regulated genes in the digestive gland that are involved in a wide variety of biological functions emphasizes the systemic response of *O. edulis* to *M. refringens*. If it remains difficult to identify a clear pattern, the identification of genes previously shown to respond to parasite infection confirms that the transcriptomic response observed may result of the *M. refringens* effects. We particularly evidenced genes involved in cilium motility or assembly (Bezares-Calderon et al., 2020) or genes encoding G-protein-coupled receptors (14 genes) that are involved in the regulation of several cellular processes by sensing molecular cues outside the cells may play a role in the immune response (He et al., 2015 ; Song et al., 2015). The identification of several genes up-regulated in infected oysters that are involved in DNA, RNA or histone methylation process suggests that epigenetic processes could participate to the response to *M. refringens*. A growing number of studies suggested that protozoan parasites such as *Leishmania, Toxoplasma*, and *Theileria* manipulate host cells *via* epigenetic modification of host gene expression mainly resulting in permanent down-regulation of host defense mechanisms to promote intracellular replication and survival of the pathogen (Mc Master et al., 2016). If the study of the regulation of methylation processes by parasites in mollusks remains poor, the case of the *M. refringens*-*O. edulis* interaction could pave the way for new research axis. Complementary analyses involving a larger number of individuals must be undertaken to better understand the entire process of response to this parasite and to identify putative genetic markers associated with *Martelia* resistance/tolerance particularly with the objective of re-establishing the culture of *O. edulis* on the foreshore.

## Conclusion

In this study, we present a high-quality genome for the flat oyster *O. edulis* a a tool for further genomic studies. The analysis of tRNA, miRNA and TEs revealed some specificities in this genome. The preliminary transcriptomic analysis made on the response of *O. edulis* to *M. refringens*, even preliminary, open new research opportunities to better understand the molecular mechanism involved in the oyster-parasite interaction actually not understood despite the highly lethal nature of this parasite. Our study provides also a valuable reference genome for further comparative genomics and population genetic analysis or to detect signature of local adaptation including genomic re-arrangement such as chromosome inversion as observed in some *O. edulis* populations (data not shown). This genome will also be used as a reference for research programs on QTLs related to physiological traits considered essential in aquaculture, particularly in the case of studies concerning a possible return to foreshore oyster culture.

## Data Archiving Statement

Raw sequence reads can be downloaded at the National Center for Biotechnology Information (NCBI) under BioProject PRJNA772088. The is available under the accession JAMSGK010000000

## Acknowledgements

This study was funded by the *Roscoff_O*.*edulis-V1* Fonds Européen pour les Affaires Maritimes et la Pêche (FEAMP) (grant code: PFEA470017FA1000016). We thank Génome Québec Centre d’Expertise et de Services for performing the genome sequencing and providing associated advice leading up to the work. We thank Dominique Marie for his help in flux cytometry, the GenomerPlatform of Roscoff for technical support and Yvon Madec, for providing oyster populations used in this study.

## Supplementary data

**Supplementary Figure 1:**
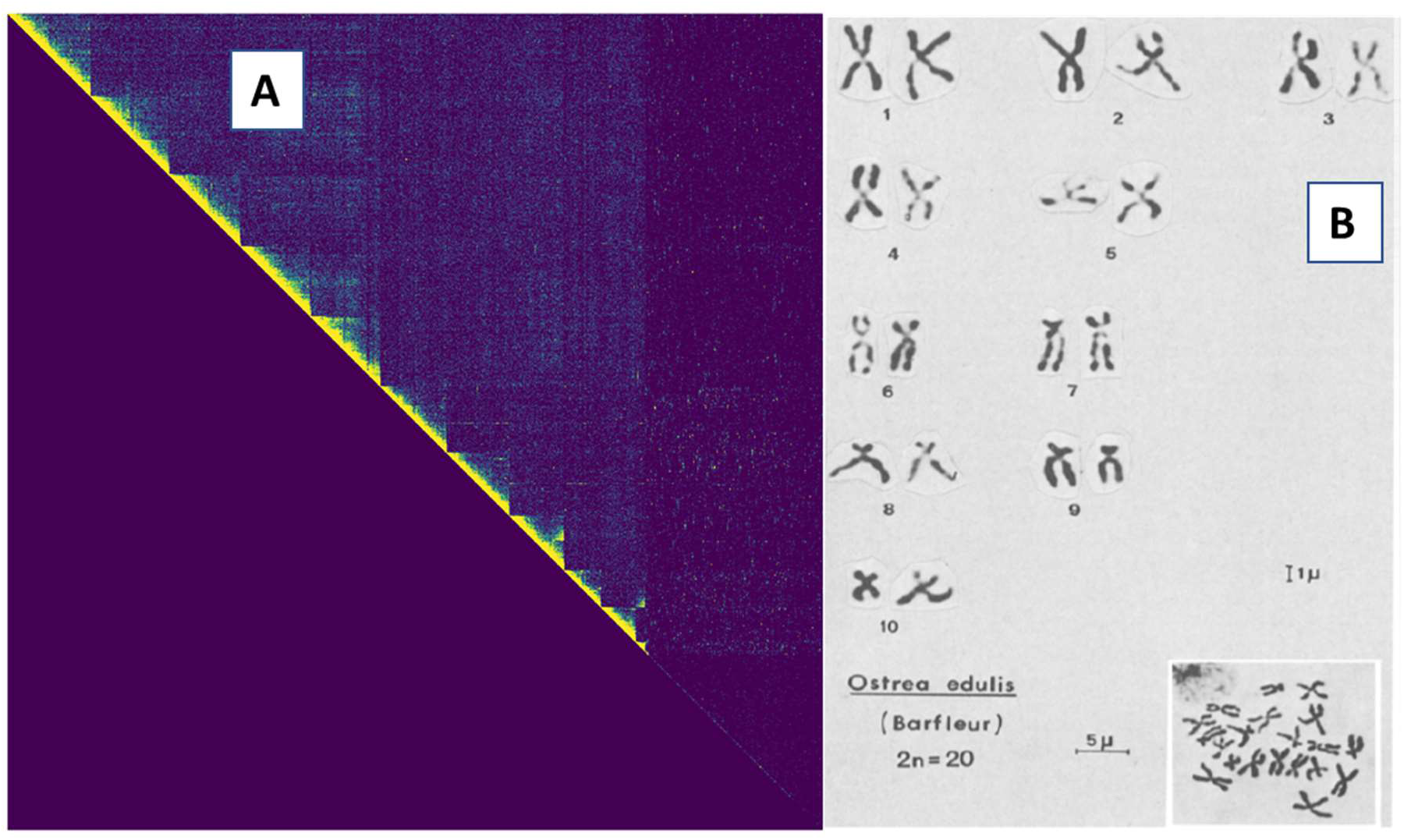
A : The 10 main scaffolds of *Roscoff_O*.*edulis-V1* genome after instaGRAAL scaffolding. B : Caryotype of *O. edulis* according to Thiriot-Quiévreux & Ayraud (1970).

**Supplementary Figure 2:**
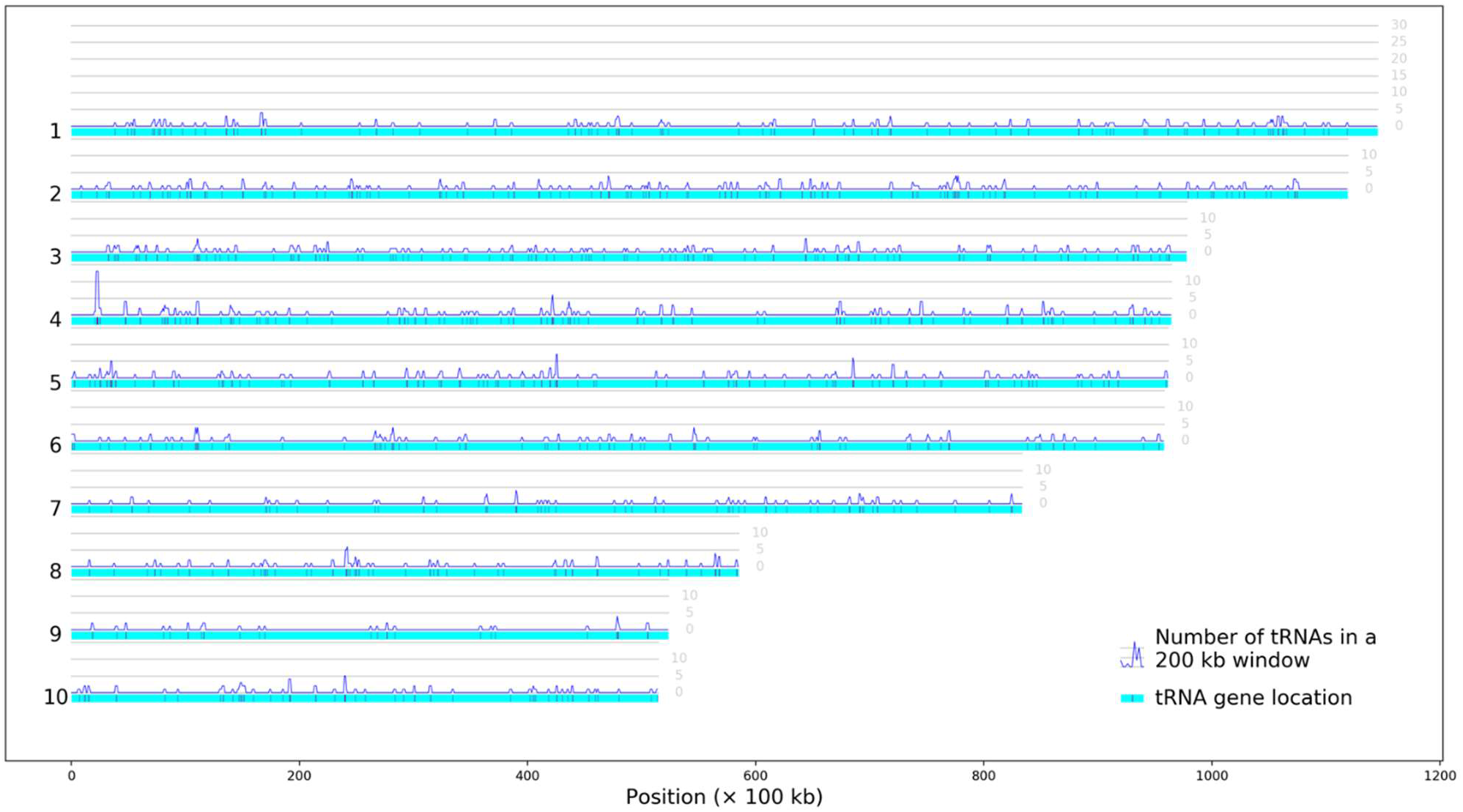
Distribution of the RNA-Ser along the 10 chromosomes in *Roscoff_O*.*edulis-V1* genome

**Supplementary Figure 3:**
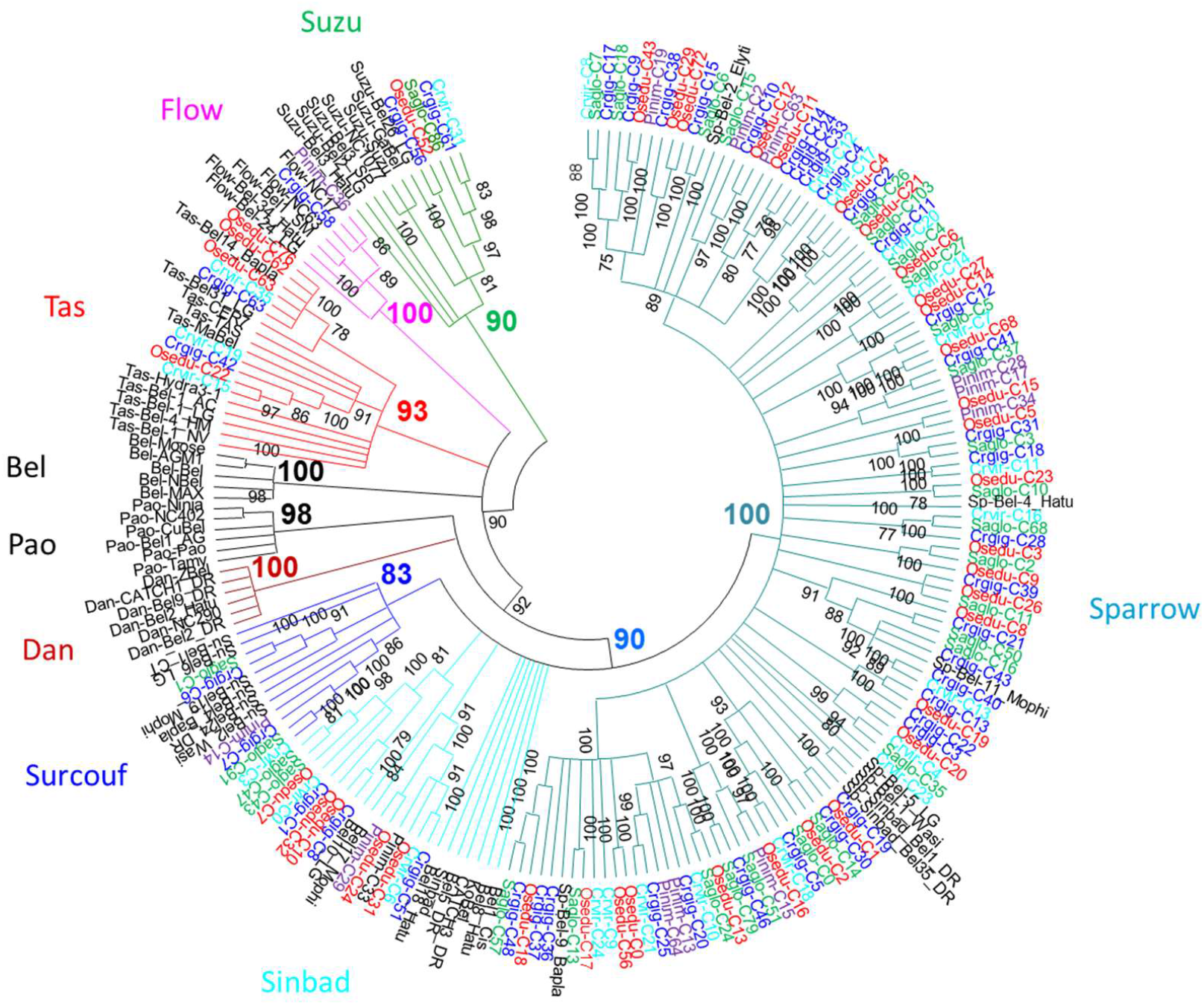
Phylogenetic relationships of BEL/Pao retrotransposons. The tree is based on Neighbor-Joining analysis of RT/RNaseH domain amino acid sequences. The BEL/Pao sub-families from oysters are indicated in color (*Crassostrea gigas* in dark blue, *Crassostrea virginica* in light blue, *Ostrea edulis* in red, *Pinctada martensii* in purple and *Saccostrea glomerata* in green) as are the clades known to possess elements of mollusks. Node statistical support values (>70 %) come from non-parametric bootstrapping using 100 replicates

**Supplementary Figure 4 :** A detailed tree representation of the clades of the Gypsy superfamily

**Supplementary Table 1** : Identification and copy number of miRNA in the genome of *O. edulis, C. virginica and M. mercenaria*. Distribution of the copy number for the 25 most represented miRNA in the 10chromosomes of the *Roscoff_O*.*edulis-V1* genome

**Supplementary Table 2 :** Genomic proportions of the sub-families and clades of LTR-retrotransposons detected in *C. gigas, C. virginica, O. edulis, P. martensii* and *S. glomerata* genomes.

**Supplementary Table 3 :** Gene ID, GOterms and enriched GOterms for Biological Process, Cellular Component and Molecular Function of genes down-regulated in *M. refringens* infected hemocytes of *O. edulis*.

**Supplementary Table 4 :** Gene ID, GOterms and enriched GOterms for Biological Process, Cellular Component and Molecular Function of genes up-regulated in Martelia infected hemocytes of *O. edulis*.

**Supplementary Table 5 :** Gene ID, GOterms and enriched GOterms for Biological Process, Cellular Component and Molecular Function of genes up-regulated in Martelia infected digestive gland of *O. edulis*.

**Supplementary Table 6 :** Gene ID, GOterms and enriched GOterms for Biological Process, Cellular Component and Molecular Function of genes down-regulated in Martelia infected digestive gland of *O. edulis*.

## References

Admire, A., Shanks, L., Danz, N., Wang, M., Weier, U., Stevens, W., … Weinert, T. (2006). Cycles of chromosome instability are associated with a fragile site and are increased by defects in DNA replication and checkpoint controls in yeast. Genes and Development, 20(2), 159–173. doi: 10.1101/gad.1392506

Altschul, S.F., Madden, T.L., Schäffer, A.A., Zhang, J., Zhang, Z., Miller, W., Lipman, D.J. (1997). Gapped BLAST and PSI-BLAST: a new generation of protein database search programs. Nucleic Acids Research, 25(17), 3389–402. doi: 10.1093/nar/25.17.3389.

Bai, C.M., Xin, L.S., Rosani, U., Wu, B., Wang, Q.C., Duan, X.K., Liu, Z.H., Wang, C.M. (2019). Chromosomal-level assembly of the blood clam, Scapharca (Anadara) broughtonii, using long sequence reads and Hi-C. Gigascience, 8(7): giz067. doi: 10.1093/gigascience/giz067

Baudry, L., Guiglielmoni, N., Marie-Nelly, H., Cormier, A., Marbouty, M., Avia, K., … Koszul, R. (2020). instaGRAAL: chromosome-level quality scaffolding of genomes using a proximity ligation-based scaffolder. Genome Biology, 21(1), 148. doi: 10.1186/s13059-020-02041-z

Beck, M.W., Brumbaugh, R.D., Airoldi, L., Carranza, A., Coen, L.D., Crawford, C., … Guo, X. (2011). Oyster Reefs at Risk and Recommendations for Conservation, Restoration, and Management. BioScience, 61, 107–116

Bennetzen, J.L. (2000). Transposable element contributions to plant gene and genome evolution. Plant Molecular Biology, 42(1), 251–269

Bermudez-Santana, C., Stephan-Otto Attolini, C., Kirsten, T., Engelhardt, J., Prohaska, S.J., Steigele, S., Stadler, P.F. 2010. Genomic organization of eukaryotic tRNAs. BMC Genomics, 11:270. http://www.biomedcentral.com/1471-2164/11/270

Bernatchez, L., Wellenreuther, M., Araneda, C., Ashton, D. T., Barth, J. M. I., Beacham, T. D., Withler, R. E. (2017). Harnessing the power of genomics to secure the future of seafood. Trends in Ecology & Evolution, 32(9), 665–680. https://doi.org/10.1016/j.tree.2017.06.010

Bolger, A.M., Lohse, M., Usadel, B. (2014). Trimmomatic: a flexible trimmer for Illumina sequence data. Bioinformatics, 30, 2114–2120. doi:10.1093/bioinformatics/btu170

Bueno, M.J., Malumbres, M. (2011). MicroRNAs and the cell cycle. Biochimica Biophysica Acta, 1812(5), 592–601. doi: 10.1016/j.bbadis.2011.02.002

Camacho, C., Coulouris, G., Avagyan, V., Ma, N., Papadopoulos, J., Bealer, K., Madden, T.L. (2009). BLAST+: Architecture and applications. BMC Bioinformatics, 10(1), 1–9. doi: 10.1186/1471-2105-10-421

Capy, P, Gasperi, G, Biemont, C, Bazin, C. (2000). Stress and transposable elements: co-evolution or useful parasites? Heredity, 85(Pt 2), 101–106. doi: 10.1046/j.1365-2540.2000.00751.x

Chan, J., Wang, L., Mu, K., Bushek, D., Xu, Y., Guo, X., … Zhang, L. (2021). Transcriptomic response to Perkinsus marinus in two Crassostrea oysters reveals evolutionary dynamics of host-parasite interactions. Frontiers in Genetics, 12, 795706. doi: 10.3389/fgene.2021.795706

Chen, G., Zhang, C., Jiang, F., Wang, Y., Xu, Z., Wang, C. (2014). Bioinformatics analysis of hemocyte miRNAs of scallop Chlamys farreri against acute viral necrobiotic virus (AVNV). Fish Shellfish Immunology,37(1), 75–86. doi: 10.1016/j.fsi.2014.01.002

Chu, Q., Sun, Y., Cui, J., Xu, T. (2017). MicroRNA-3570 Modulates the NF-κB Pathway in Teleost Fish by Targeting MyD88. Journal Immunology, 198(8), 3274–3282. doi: 10.4049/jimmunol.1602064

Cocci, P, Roncarati, A, Capriotti, M, Mosconi, G, Palermo, FA. (2020). Transcriptional alteration of gene biomarkers in hemocytes of wild Ostrea edulis with molecular evidence of infections with Bonamia spp. and/or Marteilia refringens parasites. Pathogens, 9(5), 323. doi: 10.3390/pathogens9050323.

Dobin, A., Davis, C.A., Schlesinger, F., Drenkow, J., Zaleski, C., Jha, S., … Gingeras, T.R. (2013). STAR: ultrafast universal RNA-seq aligner. Bioinformatics, 29(1), 15–21. doi: 10.1093/bioinformatics/bts635

Du, X., Fan, G., Jiao, Y., Zhang, H., Guo, X., Huang, R., … Liu, X. (2017). The pearl oyster Pinctada fucata martensii genome and multi-omic analyses provide insights into biomineralization. Gigascience, 1;6(8), 1–12. doi: 10.1093/gigascience/gix059

Duret, L. (2000). tRNA gene number and codon usage in the C. elegans genome are co-adapted for optimal translation of highly expressed genes. Trends in Genetic, 16, 287–289. doi: 10.1016/s0168-9525(00)02041-2

Emms, D.M., Kelly, S. (2019). OrthoFinder: Phylogenetic orthology inference for comparative genomics. Genome Biology, 20(1):1–14. doi: 10.1186/s13059-019-1832-y

Fang, Y., Xu, X.Y., Shen, Y., Li, J. (2020). miR-148 targets CiGadd45ba and CiGadd45bb to modulate the inflammatory response to bacterial infection in grass carp. Developmental Comparative Immunology, 106, 103611. doi: 10.1016/j.dci.2020.103611

Farhat, S., Bonnivard, E., Pales Espinosa, E., Tanguy, A., Boutet, I., Guiglielmoni, N., … Allam, B. (2022). Comparative analysis of the Mercenaria mercenaria genome provides insights into the diversity of transposable elements and immune molecules in bivalve mollusks. BMC Genomics, 23(1), 192. doi: 10.1186/s12864-021-08262-1

Flynn, J.M., Hubley, R., Goubert, C., Rosen, J., Clark, A.G., Feschotte, C., Smit, A.F. (2020). RepeatModeler2 for automated genomic discovery of transposable element families. Proceedings of the National Academy of Science USA, 117(17), 9451–9457. doi: 10.1073/pnas.1921046117

Francklyn, C.S., Minajigi, A. (2010). tRNA as an active chemical scaffold for diverse chemical transformations. FEBS Letters, 584, 366–375. DOI: 10.1016/j.febslet.2009.11.045

Gerdol, M., Greco, S., Pallavicini, A. (2019). 2019. Extensive Tandem Duplication Events Drive the Expansion of the C1q-Domain-Containing Gene Family in Bivalves. Marine Drugs, 17(10), 583. doi: 10.3390/md17100583

Gervais, O., Chollet, B., Dubreuil, C., Durante, S., Feng, C., Hénard, C., Lecadet, C., Serpin, D., Tristan, R., Arzul, I. (2019). Involvement of apoptosis in the dialogue between the parasite Bonamia ostreae and the flat oyster Ostrea edulis. Fish and Shellfish Immunology, 93, 958–964. doi: 10.1016/j.fsi.2019.08.035

Goodenbour, J.M., Pan, T. (2006). Diversity of tRNA genes in eukaryotes. Nucleic Acids Research, 34(21), 6137–6146. doi: 10.1093/nar/gkl725

Grabherr, M.G., Haas, B.J., Yassour, M., Levin, J.Z., Thompson, D.A., Amit, I., … Regev, A. (2011). Full-length transcriptome assembly from RNA-Seq data without a reference genome. Nature Biotechnology, 29(7): 644–652. doi: 10.1038/nbt.1883

Griffiths-Jones, S. (2005). Annotating non-coding RNAs with Rfam. Current Protocols Bioinformatics, 12, 12.5.1-12.5.12. doi: 10.1002/0471250953.bi1205s9.

Gundappa, M.K., Penazola, C., Regan, T., Boutet, I., Tanguy, A., Houtston, R.D, Bean, T., Macqueen, D.L. (2022). A chromosome level reference genome for a British European flat oyster (Ostrea edulis L.) Evolutionary Applications. (submitted).

Haas, B.J., Delcher, A.L., Mount, S.M., Wortman, J.R., Smith Jr, R.K., Hannick, L.I., … White, O. (2003). Improving the Arabidopsis genome annotation using maximal transcript alignment assemblies. Nucleic Acids Research, 31(19), 5654–66. doi: 10.1093/nar/gkg770

He, Y., Jouaux, A., Ford, S.E., Lelong, C., Sourdaine, P., Mathieu, M, Guo X. (2015). Transcriptome analysis reveals strong and complex antiviral response in a mollusk. Fish Shellfish Immunology, 46, 131–144. doi:10.1016/j.fsi.2015.05.023

Huang, D.W., Sherman, B.T., Lempicki, R.A. (2009). Systematic and integrative analysis of large gene lists using DAVID Bioinformatics Resources. Nature Protocols, 4(1), 44–57

Huang, S., Kang, M., Xu, A. (2017). HaploMerger2: rebuilding both haploid sub-assemblies from high-heterozygosity diploid genome assembly. Bioinformatics, 33(16), 2577–2579. doi: 10.1093/bioinformatics/btx220

Huang, S., Asaduzzaman, Md., Yoshitake, K., Kinoshita, S. (2021). Discovery and functional understanding of miRNAs in mollusks: a genome wide profiling approach. RNA Biology, doi: 10.1080/15476286.2020.1867798

Insua, A., Mendez, J. 1998. Physical Mapping and Activity of Ribosomal RNA Genes in Mussel Mytilus Galloprovincialis. Hereditas, 128(3), 189–194

Jurka, J., Kapitonov, V.V., Pavlicek, A., Klonowski, P., Kohany, O., Walichiewicz, J. (2005). Repbase Update, a database of eukaryotic repetitive elements. Cytogenetic and Genome Research, 110(1–4), 462–467. doi: 10.1159/000084979

Katoh, K., Rozewicki, J., Yamada, K.D. (2018). MAFFT online service: Multiple sequence alignment, interactive sequence choice and visualization. Briefing in Bioinformatics, 20(4), 1160–1166. doi: 10.1093/bib/bbx108

Kazazian, H.H. (2004). Mobile elements: drivers of genome evolution. Science, 303:1626–1632. doi: 10.1126/science.1089670.

Kenny, N.J., McCarthy, S.A., Dudchenko, O., James, K., Betteridge, E., Corton, C., … Williams, S.T. (2020). The gene-rich genome of the scallop Pecten maximus. Gigascience, 9(5):giaa037. doi: 10.1093/gigascience/giaa037. PMID: 32352532; PMCID: PMC7191990

Lallias, D., Beaumont, A.R., Haley, C.S., Boudry, P., Heurtebise, S., Lapègue, S. (2007). A first-generation genetic linkage map of the European flat oyster Ostrea edulis (L.) based on AFLP and microsatellite markers. Animal Genetics, 38(6), 560–568. doi: 10.1111/j.1365-2052.2007.01647.x

Lau, Y.T., Santos, B., Barbosa, M., Pales Espinosa, E., Allam, B. (2018). Regulation of apoptosis-related genes during interactions between oyster hemocytes and the alveolate parasite Perkinsus marinus. Fish and Shellfish Immunology, 83, 180–189. doi: 10.1016/j.fsi.2018.09.006

Lazar-Stefanita, L., Scolari, V.F., Mercy, G., Muller, H., Guérin, T.M., Thierry, A., … Koszul, R. (2017). Cohesins and condensins orchestrate the 4D dynamics of yeast chromosomes during the cell cycle. EMBO Journal, 36(18), 2684–2697. doi: 10.15252/embj.201797342

Le Roux, F., Audemard, C., Barnaud, A., Berthe, F. (1999). DNA probes as potential tools for the detection of Marteilia refringens. Marine Biotechnology, 1(6), 588–597. doi: 10.1007/pl00011814

Li, H, Handsaker, B, Wysoker, A, Fennell, T, Ruan, J, Homer, N, … 1000 Genome Project Data Processing Subgroup. (2009). The Sequence Alignment/Map format and SAMtools. Bioinformatics, 25(16), 2078–2079. doi: 10.1093/bioinformatics/btp352

Li, Y., Niu, D., Bai, Y., Lan, T., Peng, M., Dong, Z., Li, J. (2019). Characterization of the ScghC1q-1 gene in Sinonovacula constricta and its role in innate immune responses. Developmental and Comparative Immunology, 94, 16–21. doi: 10.1016/j.dci.2019.01.004

Li, A., Dai, H., Guo, X., Zhang, Z., Zhang, K., Wang, C., …Zhang, G. (2021). Genome of the estuarine oyster provides insights into climate impact and adaptive plasticity. Communications Biology, 4:1287.doi.org/10.1038/s42003-021-02823-6

Lieberman-Aiden, E., van Berkum, N.L., Williams, L., Imakaev, M., Ragoczy, T., Telling, A., … Dekker, J. (2009). Comprehensive mapping of long-range interactions reveals folding principles of the human genome. Science, 326(5950), 289–293. doi: 10.1126/science.1181369

Llorens, C., Futami, R., Covelli, L., Domínguez-Escribá, L., Viu, J.M., Tamarit, D., … Moya, A. (2011). The Gypsy Database (GyDB) of Mobile Genetic Elements: Release 2.0. Nucleic Acids Research, 39(Suppl 1), D70–74. doi: 10.1093/nar/gkq1061

Love, M.I., Huber, W., Anders, S. (2014). Moderated estimation of fold change and dispersion for RNA-seq data with DESeq2. Genome Biology, 15(12), 550. doi: 10.1186/s13059-014-0550-8

Lowe, T.M., Eddy, S.R. (1997). tRNAscan-SE: a program for improved detection of transfer RNA genes in genomic sequence. Nucleic Acids Research,25(5):955–64. doi: 10.1093/nar/25.5.955

Lu, Y.B., Yang, L.T., Qi, Y.P., Li, Y., Li, Z., Chen, Y.B., … Chen, L.S. (2014). Identification of boron-deficiency-responsive microRNAs in Citrus sinensis roots by Illumina sequencing. BMC Plant Biology, 14, 123. doi: 10.1186/1471-2229-14-123

McMaster, W.R., Morrison, C.J., Kobor, M.S. (2016). Epigenetics: A New Model for Intracellular Parasite–Host Cell Regulation. Trends Parasitology, 32(7) : 515–521. doi: http://dx.doi.org/10.1016/j.pt.2016.04.002

Marck, C., Grosjean, H. (2002). tRNomics: analysis of tRNA genes from 50 genomes of Eukarya, Archaea, and Bacteria reveals anticodon-sparing strategies and domain-specific features. RNA, 8(10), 1189–1232. doi: 10.1017/s1355838202022021

Martín-Gómez, L., Villalba, A., Kerkhoven, R.H., Abollo, E. (2014). Role of microRNAs in the immunity process of the flat oyster Ostrea edulis against bonamiosis. Infection Genetics Evolution, 27:40–50. doi: 10.1016/j.meegid.2014.06.026

McFarlane, R.J., Whitehall, S.K. (2009). tRNA genes in eukaryotic genome organization and reorganization. Cell Cycle, 8(19), 3102–3106. doi: 10.4161/cc.8.19.9625

Melayah, D., Bonnivard, E., Chalhoub, B., Audeon, C., Grandbastien, M.A. (2001). The mobility of the tobacco Tnt1 retrotransposon correlates with its transcriptionalactivation by fungal factors. Plant Journal, 28(2), 159–168. doi: 10.1046/j.1365-313x.2001.01141.x

Michaud, M., Cognat, V., Duchêne, A.M., Maréchal-Drouard, L. (2011). A global picture of tRNA genes in plant genomes. Plant Journal, 66(1), 80–93. doi: 10.1111/j.1365-313X.2011.04490.x

Milne, I., Lindner, D., Bayer, M., Husmeier, D., Mcguire, G., Marshall, D.F., Wright, F. (2009). TOPALi v2: A rich graphical interface for evolutionary analyses of multiple alignments on HPC clusters and multi-core desktops. Bioinformatics, 25(1), 126–127. doi: 10.1093/bioinformatics/btn575

Modak, T.H., Literman, R., Puritz, J.B., Johnson, K.M., Roberts, E.M., Proestou, D., Guo, X., Gomez-Chiarri, M., Schwartz, R. (2021). Extensive genome-wide duplications in th eastern oyster (Crassostrea virginica). Philosophical Transactions of the Royal Society B, 376(1825):20200164

Moyo, L., Ramesh, S.V., Kappagantu, M., Mitter, N., Sathuvalli, V., Pappu, H.R. (2017). The effects of potato virus Y-derived virus small interfering RNAs of three biologically distinct strains on potato (Solanum tuberosum) transcriptome. Virology Journal, 14(1):129. doi: 10.1186/s12985-017-0803-8

Mun, S., Kim, Y.J., Markkandan, K., Shin, W., Oh, S., Woo, J., … Han, K. (2017). The Whole-Genome and Transcriptome of the Manila Clam (Ruditapes philippinarum). Genome Biology and Evolution, 9(6), 1487–1498. doi: 10.1093/gbe/evx096

Nawrocki, E.P., Eddy, S.R. (2013). Infernal 1.1: 100-fold faster RNA homology searches. Bioinformatics, 29(22), 2933–5. doi: 10.1093/bioinformatics/btt509

Nischal, L., Mohsin, M., Khan, I., Kardam, H., Wadhwa, A., Abrol, Y.P., … Ahmad, A. (2014). Identification and comparative analysis of microRNAs associated with low-N tolerance in rice genotypes. PLoS One, 7(12), 1–13. doi: 10.1371/journal.pone.0050261

OSPAR (2009). Background document for Ostrea edulis and Ostrea edulis beds. (ed. O. Convention). Retrieved from https://www.ospar.org/documents?d=7183

Pardo, B.G., Álvarez-Dios, J.A., Cao, A., Ramilo, A., Gómez-Tato, A., Planas, J.V.,… Martínez, P. (2016). Construction of an Ostrea edulis database from genomic and expressed sequence tags (ESTs) obtained from Bonamia ostreae infected haemocytes: Development of an immune-enriched oligo-microarray. Fish and Shellfish Immunology, 59, 331–344. doi: 10.1016/j.fsi.2016.10.047

Peñaloza, C., Gutierrez, A.P, Eöry, L., Wang, S., Guo, X., Archibald, A.L, … Houston, R.D. (2021). A chromosome-level genome assembly for the Pacific oyster Crassostrea gigas. GigaScience, 10(3), giab020. doi: 10.1093/gigascience/giab020

Peng, J, Li, Q, Xu, L, Wei, P, He, P, Zhang, X, … Chen, X. (2020). Chromosome-level analysis of the Crassostrea hongkongensis genome reveals extensive duplication of immune-related genes in bivalves. Molecular Ecology Resources, 20(4), 980–994. doi: 10.1111/1755-0998.13157

Rocha, E.P.C. 2004. Codon usage bias from tRNA’s point of view: Redundancy, specialization, and efficient decoding for translation optimization. Genomes Research, 14:2279–2286

Ronza, P., Cao, A., Robledo, D., Gómez-Tato, A., Álvarez-Dios, J.A., Hasanuzzaman, A.F.,… Martínez, P. (2018). Long-term affected flat oyster (Ostrea edulis) haemocytes show differential gene expression profiles from naïve oysters in response to Bonamia ostreae. Genomics, 110(6), 390–398. doi: 10.1016/j.ygeno.2018.04.002

Rosani, U., Abbadi, M., Green, T.J., Bai, C.M. (2020). Parallel analysis of miRNAs and mRNAs suggests distinct regulatory networks in Crassostrea gigas infected by Ostreid herpesvirus 1. BMC Genomics 21(1), doi:10.1186/s12864-020-07026-7

Shen, Y., Yue, G. (2019). Current status of research on aquaculture genetics and genomics-information from ISGA 2018. Aquaculture and Fisheries, 4(2), 43–47

Simão, F.A., Waterhouse, R.M., Ioannidis, P., Kriventseva, E.V., Zdobnov, E.M. (2015). BUSCO: assessing genome assembly and annotation completeness with single-copy orthologs. Bioinformatics, 31(19), 3210–3212. doi: 10.1093/bioinformatics/btv351

Smaal, A.C., Kamermans, P., van der Have, T.M., Engelsma, M., Sas, H.J.W. (2015). Feasibility of Flat Oyster (Ostrea edulis L.) restoration in the Dutch part of the North Sea. Institute for Marine Resources & Ecosystem Studies, 1–58

Song, L., Wang, L., Zhang, H., Wang, M. (2015). The immune system and its modulation mechanism in scallop. Fish Shellfish Immunology, 46, 65–78. doi:10.1016/j.fsi.2015.03.013

Song, H., Guo, X., Sun, L., Wang, Q., Han, F., Wang, H., … Zhang, T. (2021). The hard clam genome reveals massive expansion and diversification of inhibitors of apoptosis in Bivalvia BMC Biology, 19(1), 15. doi: 10.1186/s12915-020-00943-9

Stanke, M., Schöffmann, O., Morgenstern, B., Waack, S. (2006). Gene prediction in eukaryotes with a generalized hidden Markov model that uses hints from external sources. BMC Bioinformatics, 7(1), 62. doi: 10.1186/1471-2105-7-62

Tamura, K., Peterson, D., Peterson, N., Stecher, G., Nei, M., Kumar, S. (2011). MEGA5: Molecular evolutionary genetics analysis using maximum likelihood, evolutionary distance, and maximum parsimony methods. Molecular Biology and Evolution, 28(10), 2731–2739. doi: 10.1093/molbev/msr121

Thiriot-Quiévreux, C., Ayraud, N. (1982). Les caryotypes de quelques espèces de bivalves et de gastéropodes marins. Marine Biology, 70, 165–172. doi: 10.1007/BF00397681

Thomas-Bulle, C., Piednoël, M., Donnart, T., Filée, J., Jollivet, D., Bonnivard, É. (2018). Mollusk genomes reveal variability in patterns of LTR-retrotransposons dynamics. BMC Genomics, 19(1), 1–18. doi: 10.1186/s12864-018-5200-1

Vera, M., Pardo, B.G., Cao, A., Vilas, R., Fernández, C., Blanco, A., … Martínez, P. (2019). Signatures of selection for bonamiosis resistance in European flat oyster (Ostrea edulis): New genomic tools for breeding programs and management of natural resources. Evolutionary Applications, 12(9), 1781–1796. doi: 10.1111/eva.12832

Wang, Q., Yu, Y., Li, F., Zhang, X., Xiang, J. (2017). Predictive ability of genomic selection models for breeding value estimation on growth traits of Pacific white shrimp Litopenaeus vannamei. Chinese Journal Oceanology. Limnology, 35, 1221–1229. doi: 10.1007/s00343-017-6038-0

Yáñez, J.M., Newman, S., Houston, R.D. (2015). Genomics in aquaculture to better understand species biology and accelerate genetic progress. Frontiers Genetics, 6, 128. doi: 10.3389/fgene.2015.00128

Yang, J. L., Feng, D.D., Liu, J., Xu, J.K., Chen, K., Li, Y.F., … Lu, Y. (2021). Chromosome-level genome assembly of the hard-shelled mussel Mytilus coruscus, a widely distributed species from the temperate areas of East Asia. GigaScience, 10(4), giab024. doi: 10.1093/gigascience/giab024

Zimin, A.V., Marçais, G., Puiu, D., Roberts, M., Salzberg, S.L., Yorke, J.A. (2013). The MaSuRCA genome assembler, Bioinformatics, 29(21), 2669–2677. https://doi.org/10.1093/bioinformatics/btt476

