## Supplementary figures and images for "Chromosomal assembly of the flat oyster (*Ostrea edulis* L.) genome as a new genetic ressource for aquaculture"

### SupplementaryFigure3

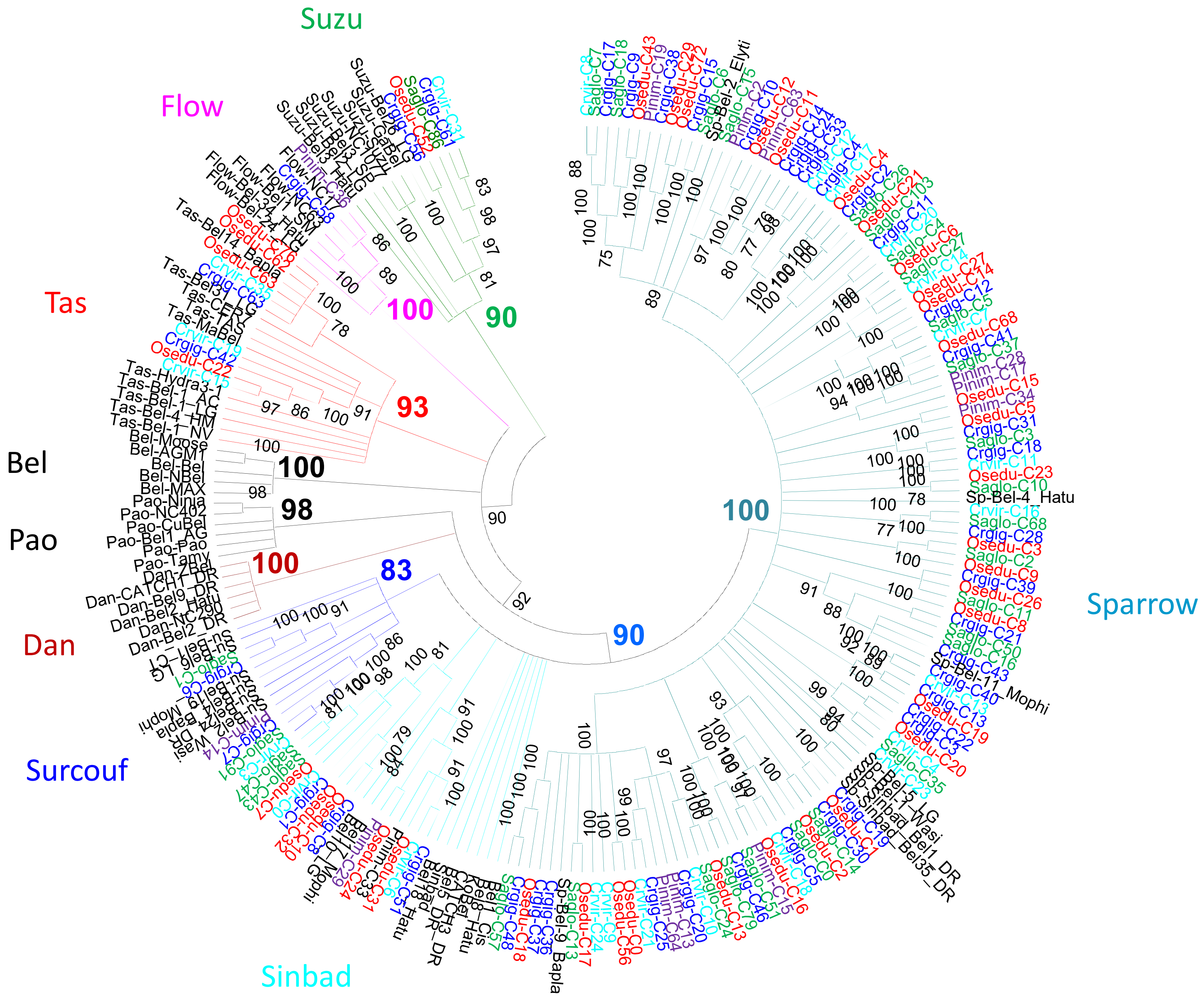
