## SupplementaryFigure4 for "Chromosomal assembly of the flat oyster (*Ostrea edulis* L.) genome as a new genetic ressource for aquaculture"

### Slide 1
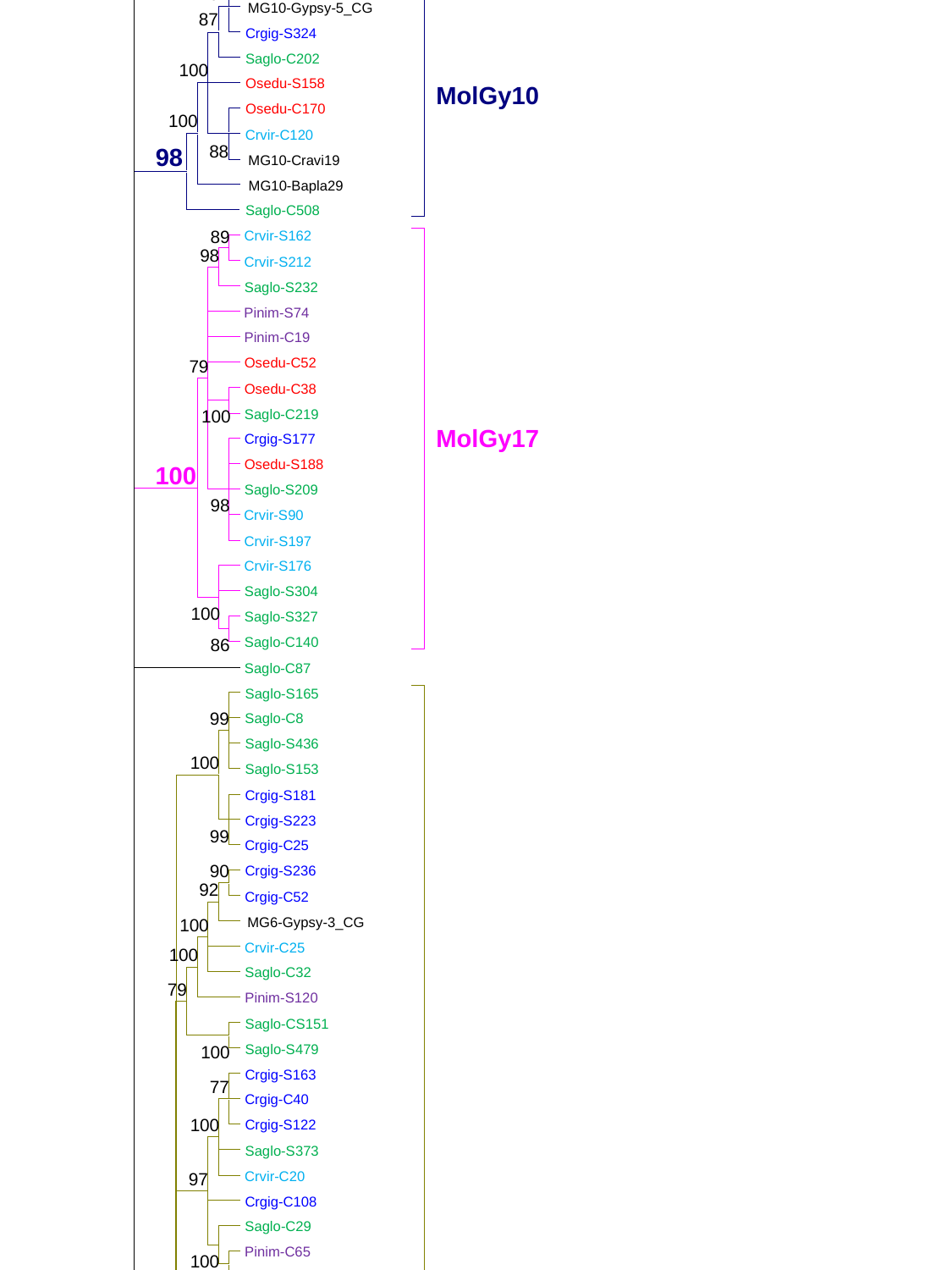

100
100
100
100
100
100
100
100
100
100
100
100
100
100
100
100
100
100
100
100
100
100
100
100
100
100
100
100
100
100
100
100
100
100
100
100
100
100
100
100
100
100
100
100
100
100
100
100
100
100
100
100
100
100
100
100
100
100
100
100
100
100
100
100
100
100
100
100
100
100
100
100
100
100
100
100
100
100
100
100
100
100
100
100
100
100
100
100
100
100
100
100
100
100
100
100
100
100
100
100
100
100
100
100
100
100
100
100
100
100
100
C-clade
100
100
100
100
 Crgig-S313
 Saglo-C287
 Osedu-S348
 Osedu-S399
 Crvir-C186
 Saglo-S388
 Saglo-S492
 Saglo-C234
 Crvir-C205
 Osedu-C318
 Crvir-S113
 Crvir-S234
 Osedu-S162
 Saglo-S459
 Pinim-C230
 Pinim-C203
 Crgig-S358
 Osedu-C285
 Saglo-S636
 Saglo-C340
 Osedu-C330
 Crgig-C280
 Saglo-S233
 Crgig-S357
 Saglo-C315
 Crvir-C217
 Osedu-S127
 Crgig-S316
 Osedu-S366
 Osedu-C46
 Osedu-S88
 Crgig-C79
 Osedu-C47
 Saglo-C42
 Saglo-S197
 Crgig-C95
 Saglo-S360
 Saglo-C144
 Crvir-C86
 Osedu-S193
 Osedu-S241
 Osedu-S202
 Osedu-S254
 Osedu-S161
 Saglo-S118
 Saglo-C119
 Crgig-S360
 Crgig-C80
 Crvir-C42
 Crvir-S64
 Pinim-C12
 Pinim-C95
 Pinim-C84
 Pinim-S333
 Crvir-C52
 Crvir-C56
 Crvir-C125
 Osedu-S219
 Osedu-C92
 Osedu-S209
 Osedu-S128
 Saglo-S295
 Crgig-C133
 Crgig-C48
 Saglo-S393
 Saglo-C45
 Osedu-C44
 Saglo-S184
 Crvir-C40
 Crgig-S199
 Crgig-C134
 Osedu-C39
 Crvir-C73
 Saglo-C41
 Saglo-S266
 Saglo-C193
 Saglo-C62
 Osedu-C35
 Crgig-S193
 Crgig-C72
 MG2-Nosub1
 Pinim-S108
 Pinim-C18
 Pinim-S86
 Crgig-C91
 Saglo-S112
 Osedu-C58
 Crvir-S182
 Crvir-C66
 Osedu-S118
 Osedu-CS333
 Osedu-C56
 Crvir-C48
 Saglo-S274
 Saglo-S511
 Saglo-C44
 Saglo-S400
 Crgig-S147
 Crgig-S300
 Crgig-C82
 Osedu-C53
 Osedu-S215
 Osedu-S152
MolGy2
 Pinim-S77
 Crgig-C0
 Saglo-S213
 Pinim-C244
 Osedu-S66
 Osedu-S97
 Osedu-C13
 Saglo-S177
 Saglo-S378
 Saglo-C13
 Osedu-C0
 Crvir-S223
 Crvir-C59
 Crgig-C12
 Saglo-S391
 Saglo-S99
 Crgig-S217
 Crgig-S219
 Crgig-C5
 Crvir-S167
 Crvir-C11
 Osedu-C5
 Saglo-S358
 Saglo-C18
 Pinim-C0
 Crvir-S240
 Crvir-S258
 Crvir-C106
 Crvir-C39
 Osedu-S183
 Osedu-S179
 Crgig-S104
 Saglo-CS269
 Osedu-S228
 Osedu-S168
 Osedu-C28
 Osedu-S121
 Saglo-C15
 Saglo-S260
 Saglo-C20
 Saglo-S276
 Saglo-S384
 Saglo-S163
 Saglo-C1
 Saglo-S212
 Saglo-S227
 Saglo-S207
 Pinim-S96
 Pinim-C117
 Crgig-C149
 Crgig-C179
 Saglo-S267
 Saglo-C143
 Crgig-C156
 Osedu-C99
 Crgig-C172
 Saglo-C190
 Saglo-S200
 Saglo-S426
 Pinim-S329
 Saglo-S285
 Saglo-S263
 Saglo-C142
 Osedu-C110
 Saglo-C152
 Saglo-CS249
 Crvir-S181
 Crvir-C115
 Crgig-C184
 Saglo-S372
 Saglo-S427
100
100
92
97
76
95
99
85
100
100
79
100
99
89
97
86
100
99
87
100
100
100
100
100
100
100
100
100
100
91
100
77
100
100
90
100
100
83
89
100
95
100
97
100
100
100
100
100
100
96
100
98
76
100
100
100
98
81
85
90
100
84
96
79
100
90
99
100
98
98
98
100
84
99
97
100
88
100
98
96
100
96
100
100
99
98
99
87
85
96
100
100
85
82
90
85
83
97
99
94
97
100
100
100
87
100
100
86
94
97
87
100
100
88
98
98
100
77
100
97
82
81
80
86
100
80
100
98
79
100
78
100
86
100
100
100
88
95
96
81
99
92
100
97
88
92
91
91
100
77
98
100
100
100
77
94
100
99
100
100
89
100
99
100
100
100
100
100
100
80
91
100
91
79
100
100
100
100
100
93
100
100
100
80
88
97
100
93
100
100
100
100
100
98
88
98
80
94
100
97
 Osedu-S220
 Osedu-C33
 Crvir-C32
 Crgig-C51
 Crgig-S140
 Crgig-C195
 Osedu-C95
 Osedu-S181
 Crvir-C36
 Saglo-S246
 Saglo-C40
 Saglo-C174
 Pinim-S136
 Pinim-CS236
 Pinim-S226
 Pinim-S256
 Pinim-CS22
 Osedu-S107
 Osedu-C43
 Osedu-C60
 Osedu-S302
 Osedu-S177
 Crgig-S251
 Crgig-C129
 Osedu-C86
 Crvir-S129
 Crvir-S144
 Crvir-C43
 Osedu-C83
 Saglo-S124
 Saglo-S221
 Saglo-C54
 Saglo-C194
 Crgig-S206
 Pinim-C25
 Pinim-S331
 Pinim-CS27
 MG2-Mophi10
 Osedu-S64
 Saglo-S239
 Crgig-C88
 Saglo-S206
 Osedu-S89
 Osedu-C36
 Crgig-C210
 Crgig-C56
 Saglo-S199
 Crvir-S126
 Crvir-C57
 Osedu-C109
 Saglo-C43
 Crgig-C98
 Osedu-S227
 Osedu-C100
 Osedu-S248
 Saglo-C16
 Saglo-S156
 Crvir-C17
 Osedu-C17
 Crgig-S39
 Crgig-C26
 Osedu-S61
 Crgig-S139
 Crvir-S50
 Pinim-S130
 Saglo-S353
 Pinim-S142
 Saglo-S417
95
84
76
100
98
97
100
83
84
94
90
99
86
95
79
99
100
100
100
100
76
79
100
99
100
100
94
100
83
95
100
78
100
85
100
77
100
99
88
100
83
99
100
97
99
78
82
76
83
92
100
100
81
88
79
88
100
98
97
80
87
100
98
94
75
94
100
100
89
82
86
97
98
100
83
89
81
100
97
85
84
97
97
100
99
100
85
91
96
80
98
82
88
75
75
100
79
86
99
100
100
79
100
88
85
77
97
99
88
99
83
78
84
91
84
100
87
86
81
86
81
100
82
81
81
100
96
88
79
87
96
78
76
84
88
100
87
99
83
77
100
85
87
81
100
96
86
100
75
99
88
90
95
94
90
98
98
100
88
80
86
85
100
96
100
95
76
85
100
84
97
86
78
100
88
99
88
89
92
99
84
89
100
96
78
80
84
98
99
82
84
77
94
89
95
100
79
89
100
89
98
97
98
93
97
99
96
99
98
97
100
99
99
100
100
97
100
100
100
99
100
100
96
93
98
100
98
92
99
100
100
99
100
97
89
99
100
94
100
100
100
92
100
100
90
100
100
100
100
100
99
99
99
100
92
100
98
93
100
99
100
92
100
100
91
97
99
96
100
98
100
95
100
100
100
97
91
100
97
92
97
95
99
98
95
93
99
89
89
98
99
100
90
92
97
93
100
100
98
100
97
100
100
100
99
100
92
93
94
99
93
100
100
100
100
97
100
89
98
90
92
100
100
99
97
98
97
91
100
95
98
100
97
97
92
96
95
99
100
95
91
100
100
100
96
100
93
92
96
100
100
97
99
100
100
93
97
97
96
89
95
94
99
93
94
99
100
94
97
100
98
95
99
100
100
96
94
100
99
96
100
97
100
91
92
97
91
100
90
93
90
92
99
96
99
94
100
100
91
99
 Osedu-S218
 Osedu-S102
 Osedu-C62
 Saglo-C171
 Saglo-CS320
 Crvir-S109
 Crgig-S97
 Crgig-C118
 Crvir-C58
 Crgig-S299
 Crgig-S132
 Crgig-C116
 Pinim-C155
 Pinim-S320
 Pinim-C38
 Pinim-S78
 Pinim-S97
 MG2-Gypsy-3_PF
 Crgig-S185
 Saglo-C30
 Osedu-C54
 Saglo-S161
 Saglo-S104
 Saglo-C48
 Saglo-S262
 Saglo-C23
 Osedu-C14
 Osedu-C70
 Crgig-C89
 Crgig-C150
 Crgig-C69
 Crgig-C249
 Saglo-S208
 Saglo-C145
 Osedu-S149
 Osedu-C79
 Osedu-S263
 Osedu-S305
 Crgig-C90
 Osedu-C55
 Crgig-C64
 Osedu-S155
 Saglo-CS148
 Osedu-S211
 Osedu-S81
 Osedu-C32
 Pinim-S229
 Pinim-C162
 Saglo-S307
 Saglo-S363
 Saglo-S352
 Saglo-S121
 Saglo-C115
 Saglo-C326
 Saglo-S150
 Saglo-S458
 Osedu-C29
 Crvir-S175
 Crvir-C34
 Crvir-S80
 Osedu-C34
 Crgig-C74
 Osedu-S132
 Saglo-C114
 Saglo-C36
 Saglo-S298
 Saglo-S278
 Saglo-S94
 Saglo-C204
 Saglo-C146
99
86
79
84
98
84
100
99
100
100
100
 Saglo-S251
 Pinim-S50
 Pinim-CS135
 Crgig-S205
 Crgig-C28
 Saglo-S103
 Saglo-C10
 MG2-Gypsy-31_LG
 Pinim-C90
 Crgig-S200
 Saglo-S149
 Saglo-S80
 Saglo-S385
 Osedu-S41
 Saglo-C31
 Pinim-S69
 Osedu-S104
 Crgig-S252
 Crgig-C146
 MG2-Ruphi1
 Osedu-S84
 Saglo-S56
 Crgig-S60
 Crgig-S152
 Osedu-C27
 Saglo-C28
 Pinim-C14
 Saglo-S162
 Osedu-S237
 Osedu-C30
 Pinim-S152
 Pinim-S373
 Crgig-S67
 Saglo-S211
 Crgig-C105
 Pinim-C13
 Saglo-C198
 Osedu-S131
 Crvir-C6
 Crgig-C20
 Saglo-S243
 Saglo-S432
 Osedu-C9
 Saglo-S167
 Crgig-S120
 Crgig-C29
 Pinim-S201
 Pinim-S45
 Saglo-S247
 Osedu-S358
 Osedu-C145
 Crgig-C23
 Osedu-C2
 Osedu-C22
 Saglo-S105
 Saglo-C0
 Osedu-S148
 Osedu-S156
 Crgig-C153
 Crgig-C50
 Crgig-C11
 Crgig-C112
 Saglo-C89
 Osedu-S159
 Crgig-S220
 Osedu-S80
 Saglo-C139
 Osedu-S142
 Osedu-C173
 MG2-Gypsy-10_BG
 Pinim-S140
 Pinim-S171
 Pinim-C4
94
83
100
94
 Pinim-S106
 Pinim-S134
 Osedu-S77
 Pinim-S110
 Crvir-S68
 Saglo-S63
 Saglo-S217
 Crgig-S87
 Crgig-C21
 Saglo-S185
 Pinim-CS100
 Pinim-CS41
 Pinim-S156
 Saglo-S123
 Crgig-S117
 Crvir-S208
 Crvir-S37
 Saglo-C69
 Pinim-C62
 Pinim-S46
 Pinim-CS31
 Pinim-CS10
 Pinim-S209
 Crvir-S163
 Osedu-C21
 Crgig-S137
 Osedu-S69
 Saglo-S79
 Saglo-C11
 Osedu-C24
 Osedu-S63
 Osedu-C26
 Crvir-S99
 Crgig-C44
 Osedu-S65
 Saglo-C64
 Saglo-C82
 Saglo-S319
 Crgig-S136
 Saglo-C84
 Crvir-C23
 Saglo-S237
 Crgig-C62
 Saglo-C38
 Saglo-S126
 Pinim-CS36
 Saglo-S74
 Saglo-C25
 Crgig-S230
 Crgig-C53
 Pinim-S139
 Saglo-C101
 Crgig-C161
 Saglo-S366
 Crgig-S165
 Crgig-C65
 Saglo-C37
 Pinim-S347
 Pinim-C17
 Crgig-S351
 Crgig-C45
 Saglo-S173
 Saglo-S187
 Pinim-S137
 Pinim-S354
 Pinim-S63
 Crgig-CS144
 Saglo-S138
 Saglo-C21
 Crgig-S293
 Osedu-S112
 Osedu-S163
87
99
97
 Crgig-S102
 Saglo-S113
 Saglo-S49
 Saglo-S313
 Saglo-S50
 Crgig-S55
 Crvir-S27
 Osedu-C1
 Pinim-S126
 Saglo-C51
 Pinim-S42
 Pinim-C93
 Saglo-C86
 Osedu-S212
 Osedu-C31
 Osedu-C98
 Pinim-S24
 Saglo-S39
 Crvir-S61
 Osedu-S59
 Crgig-S78
 Saglo-S159
 Saglo-S58
 Pinim-S111
 Osedu-S48
 Crgig-S162
 Saglo-S189
 Crgig-C167
 Crgig-C76
 Saglo-S182
 Crvir-C45
 Saglo-S256
 Pinim-C16
 Pinim-C20
 Crvir-C158
 Crvir-C67
 Saglo-S141
 Saglo-C46
 Crgig-S270
 Saglo-C117
 Crvir-S46
 Crvir-C14
 Crgig-C37
 Saglo-S155
 Saglo-S172
 Osedu-S147
 Pinim-S44
 Crgig-C47
 Crvir-S96
 Crvir-C22
 Crgig-C160
 Saglo-S188
 Crvir-C0
 Osedu-C96
 Crgig-S244
 Crgig-S94
 Crgig-S86
 Saglo-CS346
 Crgig-C57
 Crgig-S178
 Osedu-C19
 Saglo-S196
 Saglo-S259
 Crvir-S111
 Saglo-S95
 Osedu-S76
 Pinim-CS76
 Pinim-S383
 Pinim-S59
 Pinim-C1
 Pinim-C11
 Crvir-C3
 Saglo-S61
 Saglo-C100
100
94
92
100
99
100
99
89
97
97
90
MolGy1
 Saglo-S164
 Saglo-C130
 Crvir-S47
 Crvir-S121
 Osedu-S390
 Osedu-S133
 Crvir-C26
 Saglo-C17
 Saglo-C77
 Crgig-S99
 Crgig-C13
 Crgig-C66
 Crvir-S62
 Osedu-C3
 Crvir-S236
 Saglo-S439
 Crgig-C10
 Osedu-C57
 Saglo-C7
 Crgig-S338
 Crvir-C9
 Osedu-C4
 Crvir-C51
 Crvir-C8
 Crgig-S126
 Osedu-C7
 Saglo-S81
 Crgig-S103
 Crvir-S54
 Saglo-C6
 Saglo-S108
 Saglo-S410
 Crgig-C198
 MG10-Gypsy-5_CG
 Crgig-S324
 Saglo-C202
 Osedu-S158
MolGy10
 Osedu-C170
 Crvir-C120
 MG10-Cravi19
 MG10-Bapla29
 Saglo-C508
 Crvir-S162
 Crvir-S212
 Saglo-S232
 Pinim-S74
 Pinim-C19
 Osedu-C52
 Osedu-C38
 Saglo-C219
MolGy17
 Crgig-S177
 Osedu-S188
 Saglo-S209
 Crvir-S90
 Crvir-S197
 Crvir-S176
 Saglo-S304
 Saglo-S327
 Saglo-C140
 Saglo-C87
 Saglo-S165
 Saglo-C8
 Saglo-S436
 Saglo-S153
 Crgig-S181
 Crgig-S223
 Crgig-C25
 Crgig-S236
 Crgig-C52
 MG6-Gypsy-3_CG
 Crvir-C25
 Crgig-C141
 Crgig-CS232
 Saglo-S257
 Osedu-S359
 Osedu-S402
 Crvir-S132
 Crvir-S184
 Osedu-S144
 Osedu-C185
 Saglo-S97
 Crvir-C65
 Crgig-S268
 Crgig-C71
 Pinim-C56
 Crvir-S38
 Crvir-C63
 Osedu-C213
 Saglo-S299
 Saglo-S383
 Pinim-S102
 MG1-Nosub8
 Saglo-C76
 Saglo-S261
 Saglo-S305
 Crgig-C59
 Osedu-C49
 Crvir-S102
 Crvir-C53
 Saglo-C35
 Crgig-C61
 Crvir-C33
 Osedu-S328
 Saglo-C47
 Pinim-S119
 Pinim-C98
 Saglo-S619
 Saglo-CS524
 Crgig-S311
 Crgig-C188
 Osedu-C40
 Osedu-C87
 Pinim-S125
 Pinim-C26
 Crgig-C124
 Crvir-C31
 Saglo-S271
 MG1-Elyco1
 Saglo-S70
 Saglo-S107
 Crgig-S267
 Crgig-S92
 Osedu-C11
 Osedu-S361
 Osedu-S225
 Saglo-C3
 Crvir-C10
 Crvir-S195
 Crvir-S69
 Crvir-C15
 Crgig-S213
 Crgig-C38
 Saglo-C22
 Saglo-C120
 Saglo-S110
 Osedu-S368
 Osedu-S103
 Crvir-C21
 Crgig-C246
 Crgig-C49
 Saglo-C27
 Saglo-S506
 Saglo-S218
 Saglo-S181
100
100
94
76
89
89
 Pinim-CS163
 Crvir-C16
 Crvir-S214
 Crgig-C42
 Crgig-C46
 Crvir-C19
 Saglo-C26
 Osedu-S116
 Saglo-S122
 Saglo-S154
 Crvir-CS28
 Crvir-S81
 Saglo-S137
 Osedu-S157
 Crgig-S101
 Crgig-C34
 Crvir-C79
 Saglo-C34
 Osedu-C319
 Crgig-S186
 Crgig-C19
 Crgig-C32
 Crvir-S110
 Saglo-S90
 Saglo-C24
 Saglo-S136
 Saglo-S228
 Osedu-C42
 Crvir-S93
 Crvir-C145
 Crgig-C58
 Osedu-C93
 Crvir-C41
 Saglo-C289
 Crvir-S219
 Osedu-S208
 Pinim-S124
 Saglo-S129
 Crvir-C18
 Crgig-C173
 Osedu-C85
 Saglo-S192
 Saglo-S339
 MG1-Gypsy-32_LG
 Osedu-S286
 Osedu-C37
 Crgig-S245
 Osedu-S167
 Osedu-S378
 Osedu-C23
 Crvir-S159
 Crvir-C82
 Osedu-S78
 Crvir-C24
 Saglo-S96
 Osedu-S379
 Osedu-C16
 Crgig-C31
 Saglo-C14
 Saglo-S245
 MG1-Gypsy-2_MG
 Saglo-S203
 Saglo-S402
 MG1-Bapla7
 Crgig-C157
 Saglo-S116
 Osedu-C25
 Saglo-S72
 Saglo-S92
 Saglo-C2
99
94
100
100
85
 Saglo-S240
 Saglo-S168
 Saglo-C12
 Saglo-S73
 Osedu-C166
 Saglo-S93
 Crgig-S197
 Crgig-C24
 Osedu-S306
 Osedu-C8
 Pinim-S271
 Pinim-S79
 Saglo-C9
 Crgig-C17
 Crgig-S159
 Saglo-C19
 Saglo-S125
 Saglo-C66
 Crgig-C15
 Saglo-S109
 Pinim-C5
 Pinim-S72
 Saglo-S176
 Crvir-S231
 Crvir-S95
 Osedu-S207
 Osedu-C51
 Pinim-C15
 Pinim-S237
 Pinim-S70
 Pinim-CS307
 Pinim-S29
 Crvir-C7
 Saglo-S264
 Pinim-S178
 Pinim-S66
 Pinim-S321
 Pinim-S292
 Pinim-S359
 Osedu-S146
 Osedu-C15
 Saglo-C75
 Osedu-C10
 Osedu-S136
 Osedu-S426
 Crgig-C22
 Crgig-S208
 Crgig-S169
 Osedu-S113
 Saglo-S316
 Saglo-S178
 Saglo-S464
 Saglo-S483
 MG1-Mophi34
 Crgig-S215
 Pinim-C7
 Pinim-S60
 Pinim-S94
 Crvir-C2
 Saglo-S160
 Saglo-CS4
 Saglo-S520
 Pinim-S34
 Pinim-S35
 Pinim-CS353
 Crgig-C18
 Saglo-S59
 Crgig-C27
 Crvir-C101
 Crvir-C12
 Crvir-CS216
 Crvir-S264
 Crvir-S233
 Crvir-C35
 Crvir-S55
78
89
97
100
94
96
94
79
100
 Saglo-C32
 Pinim-S120
 Saglo-CS151
 Saglo-S479
 Crgig-S163
 Crgig-C40
 Crgig-S122
 Saglo-S373
 Crvir-C20
 Crgig-C108
 Saglo-C29
 Pinim-C65
 MG6-Gypsy-25_PF
 Crgig-C3
 Crvir-C13
 Crvir-S84
 MG6-Cravi20
 Crgig-S36
 Crgig-C6
 Crvir-C5
 Crgig-S81
 Crvir-S76
MolGy6
 Pinim-S54
 Pinim-C342
 Crgig-C63
 Saglo-S220
 Crgig-S85
 Crgig-S70
 Crgig-C229
 MG6-Gypsy-21_MG
 Crgig-S131
 Crgig-S83
 Crgig-S93
 Crvir-S85
 Crgig-C2
 Saglo-S469
 Pinim-S195
 Crgig-C9
 Crvir-C4
 Crgig-C4
 MG6-Gypsy-32_CG
 Crvir-S72
 Saglo-S210
 Pinim-S67
 Crgig-S158
 Saglo-S71
 Saglo-S60
 Crvir-S78
 Crgig-S68
 Pinim-S378
 Pinim-S52
 Pinim-S371
 Crvir-S107
 Saglo-S490
 Crgig-S110
 Saglo-S179
 Pinim-S51
 Pinim-S68
 Pinim-S87
 Pinim-S33
 Crgig-S41
 Saglo-C5
 Crvir-S221
 Crvir-C139
 MG4-Clili10
 MG4-Eusco4
 Saglo-S91
 Saglo-S186
 Saglo-S180
 Crgig-C30
99
100
100
94
87
 Crvir-S146
 Crgig-S330
 Saglo-C195
 MG4-Gypsy-13_LG
 Crgig-C14
 Pinim-S91
 Pinim-S160
MolGy4
 Pinim-S389
 Pinim-S216
 Pinim-S115
 Pinim-S212
 Pinim-S9
 Pinim-C6
 Pinim-C104
 MG4-Gypsy-1_PF
 Pinim-C28
 Pinim-C2
 Pinim-S113
 Pinim-S58
 Pinim-S330
 Crgig-S180
 Crgig-S318
 Crgig-C8
 Crgig-C73
 Crgig-C227
 Crgig-C35
 Saglo-S134
 Saglo-S553
 Saglo-C33
 Osedu-C6
 Osedu-S411
 Crgig-S143
 Crgig-S190
 Crgig-C75
 Crgig-S207
 Crgig-S155
 Crgig-C33
 Osvaldo
 Ulysses
 Gmr1
Gmr1
 rGmr1
 Crgig-S174
 Crgig-C239
 Pinim-C207
 Osedu-S236
MolGy18
 Osedu-C239
 Saglo-S437
 Saglo-S462
 Saglo-C238
 Saglo-S505
 Crgig-S240
 MG7-Gypsy-23_CG
 Crgig-C114
 Crgig-S43
 Saglo-S465
 Saglo-S475
 Saglo-C258
 Osedu-C231
 Saglo-S255
 Saglo-C133
 Saglo-C270
 Crgig-C187
 Crgig-C272
 Crgig-C119
 Saglo-C78
 Crgig-S138
 Saglo-S303
 Saglo-C131
 MG7-Scabro1
 Crgig-S262
 Crgig-S77
 Crgig-C121
 MG7-Cravi3
 Crvir-C75
 Crvir-S148
 MA15-Bisia6
100
96
98
96
MolAn
 Pinim-S238
 Pinim-S315
 Pinim-S316
 Saglo-C65
 Saglo-S466
 Saglo-S293
 Crgig-C204
 Crgig-S264
 Saglo-S297
 Saglo-C55
 Crgig-C115
 Crvir-C147
 Athila4-1
Athila
 Calypso
 Diaspora
 Ogre
 Tat4-1
Tat
 Cinful-1
 RIRE2
 Crvir-S97
 Crvir-S222
 Crvir-C117
 Saglo-C170
 Saglo-S294
 Crgig-C274
 Osedu-S255
 Osedu-C154
 Osedu-C223
 MG5-Gypsy-1_LS
 MG5-Hamac6
 MG5-Gypsy-o8_LG
 Crvir-S247
 Crvir-S166
 Crvir-C140
 Crvir-S142
 Crvir-C89
 MG5-Gypsy-2_PF
 Pinim-C49
 Pinim-C319
 Pinim-C344
 MG5-Gypsy-2_BG
 Saglo-C175
 Saglo-C638
 Saglo-C248
 Saglo-S147
 Saglo-S514
 Saglo-CS425
 Saglo-S615
 Saglo-S589
MolGy5
 Crgig-S259
 Osedu-S160
 Pinim-S181
 Pinim-CS317
 Pinim-S133
 Pinim-S262
 Pinim-C32
 Pinim-C30
 Pinim-S129
 Osedu-S206
 Osedu-S108
 Osedu-S68
 Pinim-S8
 Pinim-C188
 Pinim-CS148
 Pinim-S53
 Pinim-C43
 Pinim-S298
 Pinim-S260
 Pinim-C210
 Pinim-C213
 Pinim-S284
 Pinim-S368
 Pinim-S215
 Pinim-S183
 Pinim-S146
 Cer1
 PG1-Lepcla2_DN42341
PolGy1
 PG1-Harcros2_DN354
 PG1-Harfras40_DN1315
95
100
95
 MA15-Crena16
 MA16-Clili2
 MA16-Gypsy-25_BG
 Crvir-S100
 Crvir-C70
 Osedu-C297
 MA14-Cravi2
 Crgig-S214
 Crgig-C106
 Osedu-S175
 Osedu-S325
 Pinim-S304
 Saglo-C158
 MA14-Gypsy-33_CG
 Crgig-C125
 Saglo-S418
 Saglo-S637
 Saglo-S201
 Pinim-S228
 Pinim-S290
 MA12-Potan1
 MA12-Gypsy-5_CT
 MA9-Pocan1
 MA9-Villi4
 MA8-Gypsy-2_CT
 MA8-Gypsy-8_LS
 MA-Mepal140_DN14255
 MA-Terlap2_DN5627
 MA-Terlap2_DN7397
 MA-Terlap2_DN9496
 MA-Amphi2_DN4468
 MA-Mepal140_DN3523
 Crgig-C203
 Crgig-C7
 Crgig-C84
 Saglo-C157
 Crvir-C155
 Saglo-S467
 Crgig-C100
 Crvir-S130
 Crvir-C74
 Saglo-C235
 Saglo-S365
 Saglo-S612
 Saglo-S431
 Crgig-C111
 Crgig-S175
 Crvir-C114
 Crvir-C228
 Saglo-C355
 Saglo-C83
 Osedu-S214
 Crgig-S168
 Crgig-C192
 Saglo-S191
 Saglo-S387
 Crgig-S242
 Crgig-S335
 Crgig-C171
 Saglo-C53
 Saglo-S481
 Crvir-C77
 Saglo-S244
 Saglo-S362
 Saglo-C98
 Crvir-S122
 Crvir-C91
 Crvir-S271
 Crvir-S29
 Osedu-S82
 Crgig-S275
 Crgig-S290
 Crgig-S263
 Saglo-C57
 Crvir-C71
78
100
95
93
100
95
99
100
 17.6
 Tom
17.6
 Tv1
 Zam
 Yoyo
 Nomad
Gypsy
 Gypsy
 Springer
 Cigr-1
 Cigr-Gypsy-20_OB
Cigr
 Cigr-Idino10
 Cigr-Eusco15
 Cigr-Gypsy-1_BG
 PG3-Harcros2_DN853
PolGy3
 PG3-Harful2_DN4551
 PG3-Lepcla2_DN2614
 PG3-Pefu2_DN5594
 412
412
 Mdg1
 Tor4a
 MG13-Lepwil2_DN313
 MG13-Terlap2_DN20561
MolGy13
 MG13-Gypsy-39_PF
 MG13-Gypsy-46_LG
 MG13-Gypsy-96_CG
 Woot
 Skipper
 Mdg3
Mdg3
 Micropia
 Boudicca
 CsRN1
 CsRN-Gypsy-9_CT
CsRN1
Kabuki
 CsRN-Idino3
 CsRN-Sepha1
 Ty3-1
 Amn-san
 Sushi-ichi
V-clade
 Amn-ichi
 marY1
 Skippy
Pyret
 Cgret
 Pyret
 MGLR3
 Dane-1
Maggy
 Maggy
 Grasshopper
Piggy
 Pyggy
 Real
 TF1
 Cereba
CRM
 CRM
 Beetle1
 Tse3
 G-Rhodo
 Reina
Reina
 Gimli
 Gloin
 Galadriel
Galadriel
 Monkey
 Tntom1
 REM1
 Legolas
 Bagy-1
Del
 Del
 Peabody
100
Chromoviruses
Plante
 Cer2
Cer2-3
 Cer3
 Tor2
 T2-Eusco10
Tor2
 T2-Wasci8
 T2-Sepha5
 T2-Sepma15
 Tor1
 Crgig-C16
 Saglo-S463
 Saglo-S471
 Saglo-C449
 Saglo-S456
 Saglo-C334
 Crvir-S229
 Pinim-S192
 Pinim-S247
 Saglo-C330
 Osedu-S418
 Osedu-C257
 Osedu-C341
 Saglo-C343
 Pinim-S242
 Osedu-S425
 Saglo-S448
 Saglo-C345
 AB-Gypsy-13_BG
 Pinim-S264
 Pinim-S336
 Crvir-C192
 Crgig-C285
 AB-Gypsy-8_CG
 Osedu-C274
 Saglo-S452
 Crgig-C266
 Osedu-S171
 Crgig-S243
 Crgig-S368
 Crvir-S268
 Crgig-S234
 Crvir-S239
 Osedu-S364
 Crgig-C301
 Saglo-S442
 Crvir-C206
 Osedu-C293
 Crgig-C297
 Pinim-S239
 Osedu-S391
 Osedu-C314
 Saglo-S496
 AB-Gypsy-10_PF
 Pinim-S101
 Pinim-S266
 Saglo-C354
 Crvir-C269
 Crgig-C303
 AB-CFG1
 AB-PYG1
 Pinim-S310
 Osedu-S344
 AB-Hydra2-1
 Saglo-S392
 Saglo-C359
 Saglo-S633
 Crgig-S337
 Crgig-C306
 Osedu-C326
 AB-Gypsy-20_AC
 Saglo-S409
 Saglo-C422
 Crgig-S347
 Crgig-C304
AB-clade
 Saglo-C396
 Osedu-S385
 Osedu-S429
 Crgig-S362
 Osedu-C12
 AB-Mag
 AB-Cer6
 AB-Gulliver
 AB-Gypsy-3_OB
 MG11-Terlap2_DN13588
 MG11-Hatu31
MolGy11
 MG11-Elyco24
 MG11-Gypsy-10_LG
 MG11-Harcros2_DN4422
 Saglo-S111
 Saglo-C328
 Osedu-S277
 Pinim-S253
 Pinim-S190
 Pinim-S205
 Pinim-S263
 Pinim-S71
 Saglo-S404
 Saglo-C325
 Osedu-C350
 Crvir-C187
 Saglo-S461
 Osedu-C272
 Osedu-S91
 Pinim-C191
 Saglo-C333
 Osedu-S369
 Crgig-S309
 Crvir-C196
 Saglo-C310
 Osedu-C288
 Osedu-S345
 Crvir-C183
 Osedu-C268
 Osedu-S210
 Pinim-C176
 Pinim-S261
 Pinim-S234
 Crgig-C289
 MG3-Gypsy-2_CG
 Saglo-S312
 Saglo-C351
 MG3-Gypsy-15_MG
 Crgig-S148
 Pinim-C167
 Pinim-S219
 Osedu-S269
 Saglo-S438
 Saglo-S626
 Saglo-C242
 Osedu-C290
 Crgig-C282
 Saglo-S651
 Pinim-C200
 Pinim-S245
 Pinim-C277
 MG3-Gypsy-21_AC
 MG3-Gypsy-1_LG
 Saglo-S440
 Saglo-S350
 Saglo-S296
 Saglo-S648
 Osedu-C298
 Osedu-S417
 Saglo-S408
 Saglo-C377
93
99
MolGy3
 Crgig-C281
 Osedu-C317
 Crgig-C292
 Crvir-C164
 MG3-Gypsy-17_CT
 Pinim-S343
 Pinim-C246
 Osedu-C322
 Osedu-S388
 Osedu-C294
 Pinim-S252
 Saglo-C314
 Crvir-C193
 Osedu-C307
 Osedu-S296
 Osedu-C266
 Crgig-C279
 Crvir-S210
 Crvir-C154
 Saglo-C321
 Pinim-S274
 Crvir-S261
 Osedu-S262
 Osedu-S265
 Osedu-S335
 Saglo-C324
 Crvir-S194
 Crvir-S254
 Osedu-S349
 Osedu-S424
 Pinim-S340
 Crgig-S278
 Crgig-S341
 Crgig-S365
 Crgig-C296
 Osedu-S196
 Osedu-C309
 Crvir-S225
 Saglo-S455
 Saglo-C348
 Saglo-S222
 Osedu-S299
 Osedu-C316
 Pinim-C217
 Pinim-S268
 Pinim-S328
 Pinim-C198
 Saglo-C386
 Osedu-C372
 Saglo-S226
 Crvir-S224
 Saglo-S429
 Osedu-C295
 Osedu-S258
 Osedu-C280
 Pinim-CS276
 Crvir-S220
 Crvir-C211
 Crvir-C153
 C-Gypsy-1_AC
 C-Gypsy-6_LG
 Osedu-S256
 Saglo-S420
 C-Gypsy-1_MG
 Pinim-C221
 Osedu-C289
 Saglo-C230
 Saglo-S291
 Crgig-C1
 Crgig-S327
 Crvir-S119
 Osedu-S392
 Saglo-S519
 Pinim-C154
 Saglo-S216
 Saglo-S476
98
80
89
93
86
100
96
77
91
91
99
97
 Saglo-C231
 C-Gypsy-5_LS
 Saglo-C344
 Saglo-S411
 Crgig-S325
 Osedu-C137
 Pinim-C197
 Saglo-S577
 Saglo-C369
 C-SURL
 Osedu-S190
 Saglo-S480
 Saglo-S614
 Crgig-S322
 Saglo-S485
 Pinim-C179
 Osedu-C282
 Saglo-C371
 Crgig-C320
 Pinim-C3
 Saglo-S254
 Saglo-S311
 C-Gypsy-9_CG
 Crgig-C276
 Osedu-S450
 Osedu-C278
 Saglo-S489
 Crvir-S124
 Crvir-C204
 Osedu-C300
 Saglo-S430
 Saglo-C338
 Crgig-S154
 Crgig-C310
 Saglo-C370
 Crvir-C209
 Saglo-S484
 Pinim-C220
 Pinim-C204
 Pinim-C206
 Osedu-S442
 Osedu-C281
 Crvir-C200
 Saglo-C368
 Crgig-S340
 Osedu-S153
 Osedu-C334
 Crgig-S326
 Saglo-S494
 Crvir-C199
 Osedu-S125
 Crvir-S279
 Crvir-C201
 Saglo-S445
 Osedu-C315
 Saglo-S379
 Saglo-S478
 Crgig-S314
 Saglo-S135
 Osedu-C134
 Saglo-C283
 Crvir-C190
 Crgig-S277
 Crgig-S182
 Pinim-S324
 Osedu-C284
 Crvir-S237
 Saglo-S435
 Crgig-C312
 Saglo-S398
 Osedu-C312
 Crvir-C207
 Pinim-C202
92
100
 Crgig-S258
 Crgig-C308
 Saglo-C382
 Osedu-C332
 Crvir-C215
 Crvir-S127
 Osedu-S273
 Osedu-C291
 Saglo-S503
 Saglo-S543
 Osedu-S396
 Osedu-C304
 Crgig-C288
 Crgig-C287
 Crgig-S291
 Crgig-S254
 Crvir-C202
 Saglo-S335
 Saglo-C357
 Crgig-S323
 Saglo-C356
 Osedu-C308
 Crvir-S174
 Pinim-S187
 Pinim-S381
 Pinim-S327
 Pinim-S362
 Crgig-S346
 Saglo-S428
 Osedu-C301
 Crgig-S257
 Crgig-C294
 Osedu-S124
 Osedu-C313
 Saglo-S272
 Osedu-C279
 Pinim-C177
 Crvir-S123
 Crvir-C188
 Pinim-S118
 Saglo-S401
 Crvir-C191
 Osedu-S352
 Crvir-S30
 Osedu-S393
 Crvir-S49
 Pinim-S308
 Pinim-C267
 Pinim-S149
 Crgig-C269
 Saglo-S454
 Saglo-S423
 Crgig-S336
 Saglo-C309
 Saglo-S361
 Saglo-S414
 Saglo-S486
 Osedu-C287
 Saglo-S450
 Crgig-S253
 Pinim-S243
 Crvir-S278
 Saglo-C331
 Crvir-C171
 Saglo-CS595
 Crvir-C179
 Saglo-C284
 C-Gypsy-68_CG
 Saglo-C273
 Pinim-S231
 Crgig-C237
 Saglo-C318
 Pinim-C175
 Osedu-S395
 Osedu-C283
98
100
93
100
77
100
98
75
